# Integrative identification of non-coding regulatory regions driving metastatic prostate cancer

**DOI:** 10.1101/2023.04.14.535921

**Authors:** Brian J Woo, Ruhollah Moussavi-Baygi, Heather Karner, Mehran Karimzadeh, Kristle Garcia, Tanvi Joshi, Keyi Yin, Albertas Navickas, Luke A. Gilbert, Bo Wang, Hosseinali Asgharian, Felix Y. Feng, Hani Goodarzi

## Abstract

Large-scale sequencing efforts of thousands of tumor samples have been undertaken to understand the mutational landscape of the coding genome. However, the vast majority of germline and somatic variants occur within non-coding portions of the genome. These genomic regions do not directly encode for specific proteins, but can play key roles in cancer progression, for example by driving aberrant gene expression control. Here, we designed an integrative computational and experimental framework to identify recurrently mutated non-coding regulatory regions that drive tumor progression. Application of this approach to whole-genome sequencing (WGS) data from a large cohort of metastatic castration-resistant prostate cancer (mCRPC) revealed a large set of recurrently mutated regions. We used (i) *in silico* prioritization of functional non-coding mutations, (ii) massively parallel reporter assays, and (iii) *in vivo* CRISPR-interference (CRISPRi) screens in xenografted mice to systematically identify and validate driver regulatory regions that drive mCRPC. We discovered that one of these enhancer regions, GH22I030351, acts on a bidirectional promoter to simultaneously modulate expression of U2-associated splicing factor SF3A1 and chromosomal protein CCDC157. We found that both SF3A1 and CCDC157 are promoters of tumor growth in xenograft models of prostate cancer. We nominated a number of transcription factors, including SOX6, to be responsible for higher expression of SF3A1 and CCDC157. Collectively, we have established and confirmed an integrative computational and experimental approach that enables the systematic detection of non-coding regulatory regions that drive the progression of human cancers.

## Introduction

Non-coding DNA regions are increasingly recognized as cancer drivers (1–3). However, several challenges have limited our ability to systematically annotate oncogenic non-coding genomic elements. First, for the coding genome, the recurrence of functional mutations has long been leveraged to identify cancer relevant genes (4–6). However, the paucity of whole-genome sequencing data relative to exome sequencing data limits the number of times mutations in non-coding DNA regions may be observed. This is further compounded by the much larger non-coding space relative to that of coding sequences. Secondly, while a number of heuristics have been developed to identify functional mutations in the coding genome (e.g. the ability to distinguish between sense, missense, and nonsense mutations), the concept of functionality in the non-coding space is more difficult to capture (7–12). Currently, the standard statistical approach to identify mutational hotspots in the non-coding space is to form a background distribution and use an appropriate set of covariates to detect mutational events that occur more than expected by chance above background (3,13–16). More recently, machine learning algorithms have been used to identify driver events in non-coding regions (17–20).

Nevertheless, we are not aware of any study that integrates statistical techniques using single-base-resolution machine learning platforms with state-of-the-art experimental approaches to functionally capture non-coding drivers of tumor progression. To address this gap, we developed an ensemble of statistical and deep learning models, trained on metastatic castration-resistant prostate cancer (mCRPC) genomes, to identify non-coding regulatory regions that drive prostate cancer progression. For this, we relied on whole-genome sequencing (WGS) and matched RNA-seq data generated from our recent multi-institutional study on more than 100 mCRPC patients (21). Given the genetic heterogeneity and long-tail nature of driver mutations in mCRPC (22-23), using data from a large multi-institutional study is essential to effectively capture driver regulatory elements. We then also used data generated from two separate experimental modalities to assess the functional impact of our computationally nominated regulatory elements on gene expression and tumor growth. First, we devised a massively parallel reporter assay (MPRA) to assess the impact of each mCRPC-associated region on transcriptional control (24). In parallel, we leveraged CRISPR-interference, which uses dCas9-KRAB to precisely silence chromatin at sites of interest (25), to carry out a pooled genetic screening strategy in mouse xenograft models.

By integrating data from various modules in our combined computational and experimental platform, we identified a recurrently mutated regulatory region, previously annotated as GH22I030351, that controls a bi-directional promoter driving the expression of both SF3A1, a U2-associated splicing factor, and CCDC157, a poorly characterized putative chromosomal protein. We confirmed that silencing this regulatory region in prostate cancer cell lines with CRISPRi reduced subcutaneous tumor growth. Our follow-up functional studies revealed that both SF3A1 and CCDC157 promote prostate cancer tumor growth in xenograft models. We also performed CLIP-seq and RNA-seq in SF3A1-over-expressing cells and found up-regulation to be linked to changes in the mCRPC splicing landscape. Finally, we identified multiple transcription factors, namely SOX6, that regulate expression of SF3A1 and CCDC157 upstream of GH22I030351, and functionally validated SOX6 *in vivo*, observing increased tumor growth in xenografted mice injected with SOX6-knockdown cells.

## Results

### Identifying hotspots in non-coding regions using a regression-based model

In human cancers, recurrent mutations are rare, particularly in non-coding regions of the genome. In the coding sequence space, commonly used tools such as MutSigCV (11) have been developed to assess the accumulation of mutations along the entire gene body in a given cohort to boost signal from observed mutations. We took a similar approach in the non-coding sequence space by combining counts across annotated regulatory regions in order to identify those that were recurrently mutated in our cohort of 101 mCRPC samples. However, mutational density is highly varied and heterogeneous across the genome, and broadly impacted by genetic and epigenetic factors. Therefore, to identify regulatory regions that are mutated more than expected by chance, we first needed to generate an accurate model of background mutation rates for all regions of interest.

For this, we made two key assumptions: (i) the vast majority of the non-coding regulatory regions do not harbor driver mutations and therefore are not recurrently mutated significantly above background (**Fig. S1a**), and (ii) regulatory regions with similar sequence and epigenetic features are more likely to have similar mutational densities. Given these two priors, the expected mutational density of a given region can be calculated using a predictive model trained on our cohort’s whole-genome sequencing data. Should such a model achieve high accuracy across genomic regions, its predictions can be used as a baseline estimate for expected background mutational density and can in turn be leveraged to identify significant outliers as mutational hotspots.

Since this problem is a regression analysis at its core, we took advantage of generalized linear models (GLM) to estimate mutational density in each regulatory region as a function of i) the region’s putative functional annotation, ii) sequence context, and iii) epigenetic features associated with the region, which are known to impact local mutation rates (26-27). To achieve this, we first one-hot encoded the annotated regulatory elements, generating a total of 728,208 non-overlapping genomic functional regions that were uniquely tagged (**Figs. S1b-c,** see Methods for more details). This prevented heterogeneous functional annotations within a contiguous region and ensured that each mutation in the cohort would only be counted once even if it occurred in overlapping segments. Next, to capture the sequence context, we measured dinucleotide frequencies, which are known to be non-randomly distributed. However, since the 16 dinucleotides are not entirely independent and show collinearities, we performed principal component analysis (PCA) and chose the first seven principal components, which together captured ∼80% of the total variance. Finally, as we did not have access to epigenetic data for patients in our cohort, we used the ENCODE database and picked epigenetic factors from the PC3 prostate cancer cell line as input features (covariates) to our regression model (**Table S1**). Similar to sequence context, since many of these measurements were collinear, we used a 10-PC projection of the data to represent ∼80% of variance in epigenetic space. Specifically, we used three sets of covariates: (i) a functional classification of each region, (ii) a PCA embedding of dinucleotide frequencies, and (iii) a PCA embedding of epigenetic signals (**Figs. 1a, S1d**).

**Figure 1.**
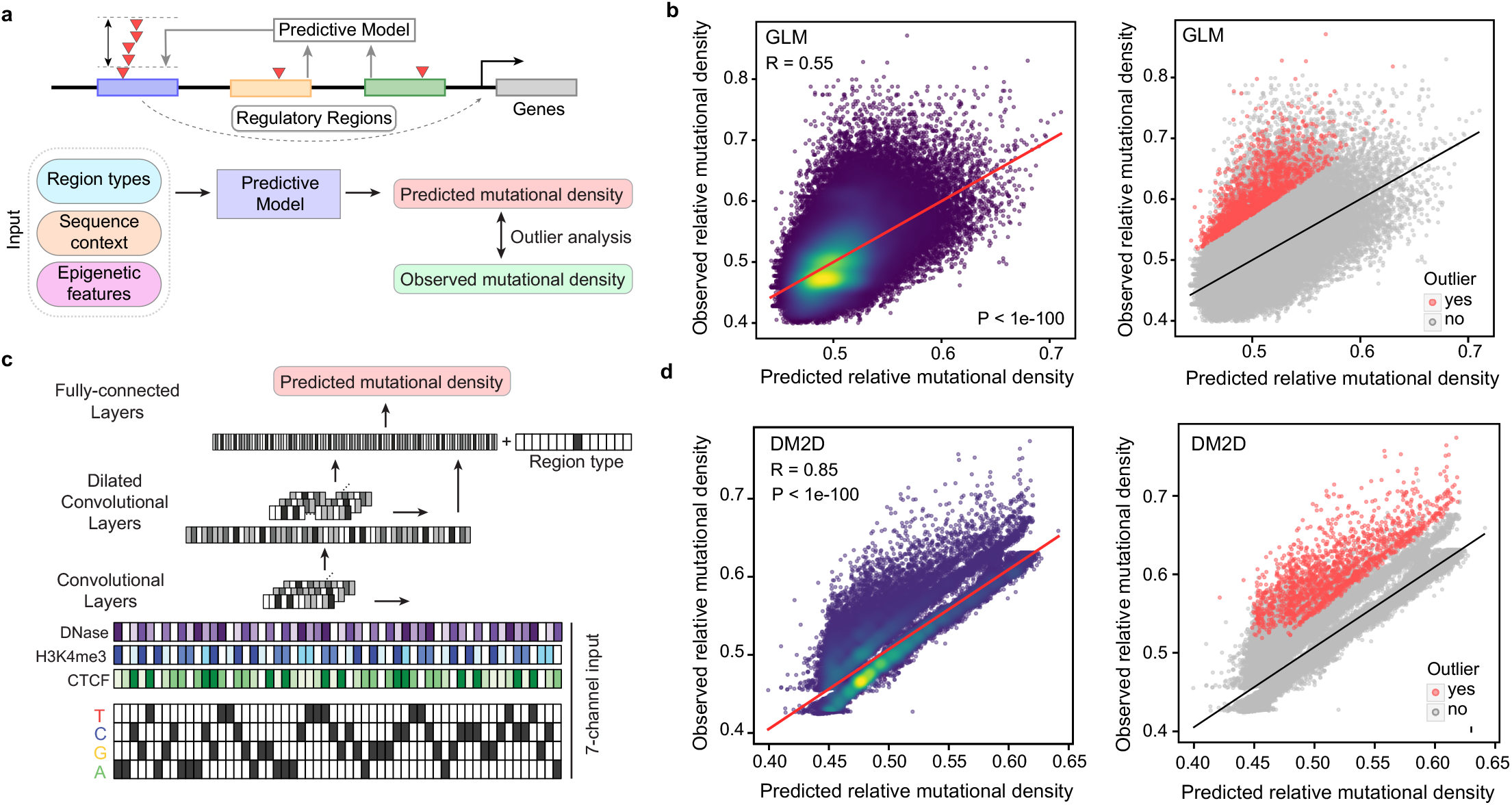
Regression and deep learning models effectively predict background mutational density in regulatory regions. **(a)** Genomic regions have a background mutation rate which is a function of their sequence context, functional annotation classes and underlying epigenetic features. We developed an outlier detection model based on a generalized linear regression model (GLM), termed MutSpotterCV, to use such features to estimate the expected mutational density in a given region. **(b)** The scatter plot of observed vs. predicted mutational density values (normalized) generated by the MutSpotterCV achieved a Pearson correlation of 0.55. We used the predictions of this model to perform an outlier analysis to identify regulatory regions that are mutated at a substantially higher rate than expected by chance. The resulting outlier regions are marked in red. **(c)** We also tested the ability of models with increased complexity to perform this prediction task. One of our best-performing models was a deep convolutional neural network (CNN). The input to this model is a multi-channel encoding of sequence and epigenetic signals. **(d)** This model, named DM2D, achieved a Pearson correlation of 0.85, far exceeding that of MutSpotterCV. Nevertheless, the identity of final outliers identified by both models were virtually the same. Therefore, we deemed these regions as regulatory elements that are hypermutated in mCRPC samples. The same outliers are colored in **(b)** and **(d)**.

We defined genomic functional regions by compiling coding- and non-coding genomic annotations–namely promoters, enhancers, promoter/enhancers, 3’UTRs, 5’UTRs, CpG islands, and gene bodies (both upstream and downstream of all annotated genes). Binary variables were created to record the affiliation of the non-overlapping genomic regions with each of the functional classes. We then mapped more than 1.8×10^6^ high-confidence, single-nucleotide variations (SNVs) and short indels present in our cohort onto these functional regions. About one in five regions had at least one mutation from at least one patient (**Fig. S1a**). Unmutated regions were excluded from the rest of the analysis. The overall average mutation frequency (mutations per Mb) in functional regions was 4.1/Mb, consistent with other whole-genome mCRPC findings (28). However, we found that mutational frequencies tended to be higher in shorter CpG islands (median: 4.91/Mb) and promoters (median: 5.60/Mb) than in longer exonic regions (median: 0.78/Mb), suggesting that observed mutations are distributed non-randomly and disproportionately with regions’ sequence length. This confirms that mutations are not uniformly distributed among functional regions, further supporting our choice to include ‘functional classes’ as a categorical covariate in our model.

We then fit a GLM-based model using mutational density as the response variable and the set of covariates described above. The resulting model, named MutSpotterCV (**M**utational density **S**potter using **C**o**V**ariates), achieved a Pearson correlation of 0.55 between observed vs. predicted mutational densities across genomic regions (**Fig. 1b**). Using MutSpotterCV, we observed a small subpopulation of regulatory regions with substantially higher observed mutational densities above that expected by chance. By systematically performing outlier detection analysis, MutSpotterCV flagged a total of 1,780 regions as a set of candidate functional regions harboring mutational hotspots (**Figs. 1b, Table S2**, see Methods for detection criteria), which amounted to 1.1% of all mutated regulatory regions. Furthermore, we found all covariates to be significantly associated with the response variable in the model, suggesting they independently and significantly contributed to the prediction of mutational density (**Fig. S1d**). In our previous study, we had identified patients in our cohort with pathogenic mutations in prostate cancer driver genes (see Table S5 in (21)). Here, we observed that a number of our non-coding mutational hotspots were proximal to a subset of prostate cancer driver genes, i.e. *AR*, *FOXA1*, and *TP53*. We therefore asked whether any of these non-coding mutational outliers were more or less likely to occur in patients with known pathogenic mutations in coding regions of these driver genes. Interestingly, we did not find any association between these non-coding mutational hotspots and pathogenic mutations in prostate cancer driver genes (*P* =0.39, two-sided Fisher’s exact test). In addition, among the 1,780 mutational hotspots identified here, six of them were found to harbor non-coding driver hits in myeloid MPN, melanoma, and prostate adenocarcinoma by a recent pan-cancer study on non-coding regions (**Table S2,** (13)). Lastly, in order to confirm the robustness of our study, we also examined the consequence of different modifications to MutSpotterCV to assess the impact of varying covariate choices on the final results (**Figs. S1e-j,** Methods).

### A multimodal convolutional neural network for accurate prediction of mutational density

We set a high threshold for detection of outliers by MutSpotterCV; however, we recognized that MutSpotterCV calls may still be dependent on the assumptions of our underlying model. Specifically, a GLM measures the linear dependence of the response variable on its predictors. Therefore, to ensure the robustness and reproducibility of our findings and capture potential nonlinear relationships among variables, we also developed a separate deep-learning-based model, termed DM2D (**D**eep **M**odel for **M**utational **D**ensity), to assess (i) whether it would be capable of achieving higher accuracy for predicting mutational density than MutSpotterCV, and (ii) the overlap between called putative mutational hotspots. DM2D is a convolutional neural network (CNN) model, which uses sequence and epigenetic data as multi-channel input with single-base resolution. As shown in **Fig. 1c**, we used a seven-channel input layer: four channels were used for one-hot encoding DNA sequence, and to ensure our results were not dependent on the choice of specifically PC3 as our prostate cancer cell line model, the other three channels were used for epigenetic data from LNCaP–namely, DNase hypersensitivity, H3K4me3 signal, and CTCF binding sites (ENCODE database). After the convolutional blocks, the resulting sequence and chromatin data embedding is combined with the functional category of the input region and passed on to a fully connected layer for mutational density prediction. Once trained, this CNN model performed substantially better than GLM, and achieved a Pearson correlation of 0.85 between observed and predicted values (**Figs. 1d, S1k**). However, this increase in accuracy was not accompanied by a significant change in identity of previously called outliers. About 90% of non-coding mutational hotspots that were detected by MutSpotterCV were also called by DM2D (**Table S2**, **Fig. S1l**).

### Quantifying the regulatory functions of identified non-coding mutational hotspots

Our focus on annotated non-coding regions was based on the underlying assumption that these regions carry out regulatory functions in gene expression control, which in turn may play a role in driving prostate cancer progression. To test this assumption, we used transcriptomic data across all patients to assess the putative effects of mutations in our non-coding mutational hotspots on gene expression. Specifically, we asked whether genes in the vicinity (within 15 kb) of these regions were significantly up- or down-regulated in the tumors that harbored mutations in cognate regions. In total, we performed differential gene expression analysis for 1,692 flanking genes in the vicinity of non-coding mutational hotspots.

For each non-coding mutational hotspot, the cohort was divided into two groups: mutant and reference. We defined mutants as patients carrying mutations in that specific hotspot and references as those that do not. We required each non-coding hotspot to include at least four patients in the mutant category, and hotspots that did not satisfy this criterion were removed (**Fig. S1m**). Using DESeq2 (29), we compared the expression of genes flanking each non-coding mutational hotspot in mutant vs. reference samples. We found 104 differentially expressed genes in the vicinity of 98 hotspots (*P* <0.05, **Tables S3-S4**; see Methods for details on selection criteria). These 98 hotspots, which we termed Candidate Driver Regulatory Regions (CDRRs), harbored a total of 885 mutations. The distribution of these mutations among tumors was scattered (**Fig. S1n**), suggesting the final CDRRs were not overly biased by a particular tumor. We noted that one of our CDRRs, located in the 3’ UTR of the oncogene *FOXA1*, was also identified as a non-coding driver in prostate cancer by a recent pan-cancer study (13).

Next, to functionally validate these CDRRs, we used a massively parallel reporter assay (MPRA), which allows for scalable measurement of enhancer activity across thousands of sequences (30). As schematized in **Fig. 2a**, MPRA analysis involves measuring the difference between enhancer activity associated with each fragment and its matched scrambled control. This activity is calculated by comparing the ratio of reference/scrambled in the RNA population to the same ratio in genomic DNA (gDNA) samples, which captures their representation in the original library. In our MPRA analysis, performed in biological triplicate (**Fig. S2a**), barcodes assigned to 358 fragments of interest and their scrambled controls were observed at sufficient read counts for downstream analyses (>25 reads per barcode). Specifically, we included in our MPRA library the reference human genome sequences for each fragment, as well as all mutant variants observed in our patient cohort. We used logistic regression to compare enhancer activity between reference and scrambled sequences. At an FDR of <0.01 and effect size of 1.5-fold differential expression, roughly a third of our fragments showed a significant effect on transcriptional activity (**Fig. 2b**).

**Figure 2.**
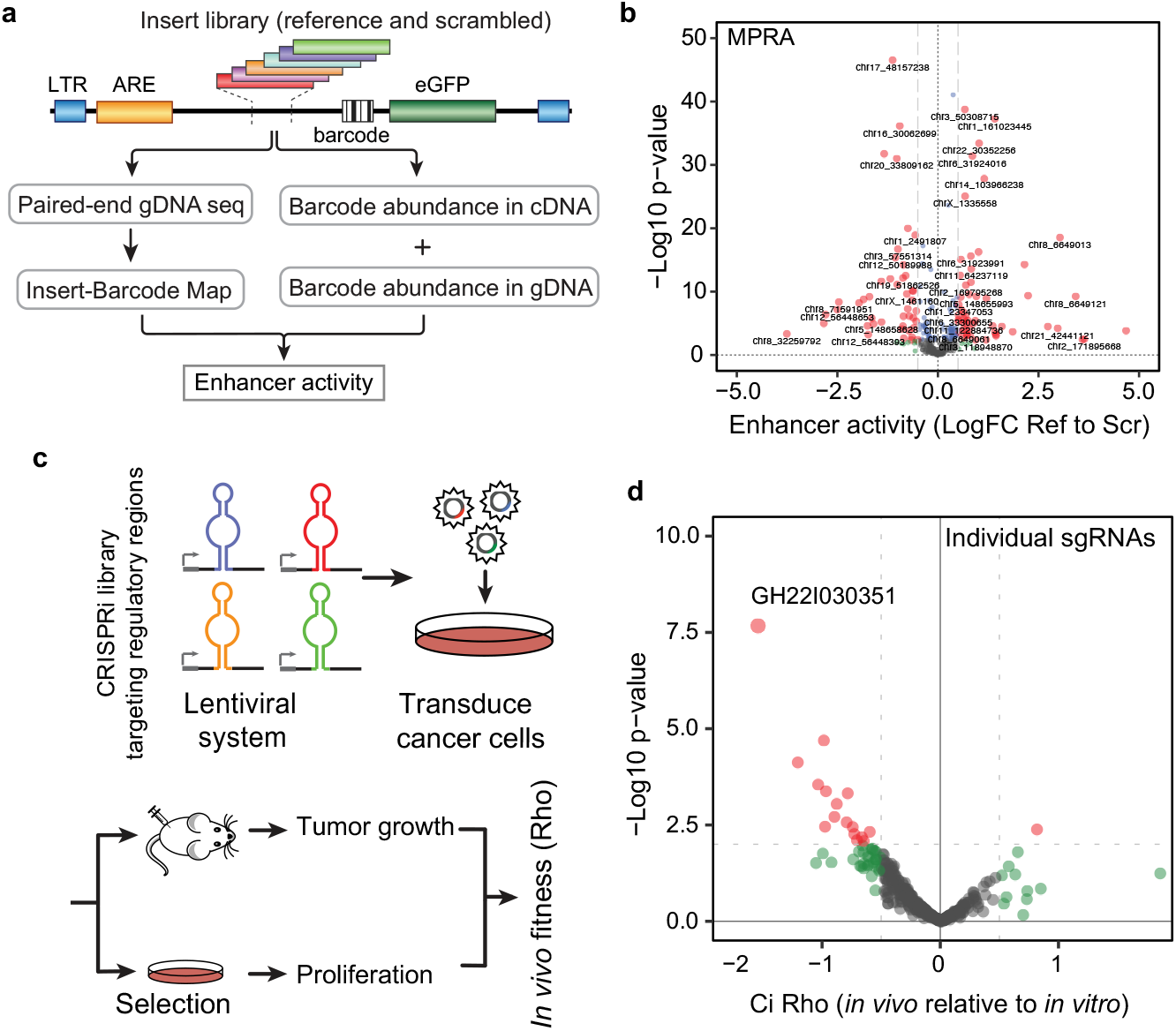
Regulatory and fitness consequences of mCRPC-associated non-coding regulatory regions. **(a)** The schematic of the massively parallel reporter assay (MPRA) used to assess the enhancer activity of regulatory sequences hyper-mutated in mCRPC and their scrambled control as background. **(b)** A volcano plot showing the measured enhancer activity for each regulatory segment (WT sequence) relative to its scrambled control. **(c)** Schematic of our *in vivo* CRISPRi strategy designed to identify regulatory regions that contribute to subcutaneous tumor growth in xenografted mice. **(d)** *In vivo* fitness consequences of expressing sgRNAs targeting mCRPC hyper-mutated regulatory regions. The x-axis shows the calculated fitness scores (Rho), where positive values denote increased tumor growth upon sgRNA expression and negative values denote the opposite. The y-axis represents -log10 of *P*-value associated with each enrichment.

In order to reveal potential *cis*-regulatory elements that are embedded in these functionally active regions, we performed regulon analysis as well as *de novo* motif discovery. We asked whether there were binding sites associated with any known transcription factors that were significantly enriched among the regions with regulatory activity in our MPRA system. For this, we systematically intersected annotated binding sites (narrowPeaks) from the ENCODE database across all profiled transcription factors with the population of fragments cloned in our MPRA library. We then used iPAGE (31) to ask whether these annotated binding sites showed a significant association with enhancer activity. This analysis revealed JunD, an AP-1 transcription factor, to be significantly associated with increased enhancer activity in our MPRA system (**Fig. S2b**). This is consistent with the known role of AP-1 factors as foundational drivers of prostate cancer progression (32-33). For example, it has been shown that JunD has an essential role in prostate cancer cell proliferation, and also is a key regulator for cell cycle-associated genes (34). JunD employs c-MYC signaling to regulate prostate cancer progression, and is a coactivator for androgen-induced oxidative stress–a key role player in the prostate cancer onset and progression (35-36). In addition to the analysis described above, which relies on annotated binding sites, we also used the primary sequence of our fragments to directly perform *de novo* motif discovery using FIRE (37). As shown in **Fig. S2b**, we discovered two motifs, one of which has similarities to the binding site of the transcription factor SMAD. Overall, the MPRA analysis revealed fragment-level readouts of transcriptional activity, and the putative regulators that underlie their activity.

Given that for the majority of putative regulatory regions more than one fragment per mutation was included in our MPRA library, we then performed a region-level analysis by integrating measurements for the fragments across each region. Achieving statistical significance in this analysis would require concordant effects from multiple fragments in the same direction, highlighting the functional relevance of the identified regulatory regions and providing a rational approach for prioritizing their collective impact on gene expression (**Fig. S2c**). Taken together, results from our endogenously controlled MPRA highlights the strong enrichment of regulatory sequences in CDRRs associated with mCRPC.

### A systematic CRISPR-interference screen for non-coding drivers in xenograft models

Our analyses of gene expression data from mutated and unmutated samples for each region of interest, coupled with a large-scale and systematic MPRA analysis, provided strong evidence for many of our CDRRs to have a regulatory function in gene expression control. However, to be considered an actual driver of prostate cancer progression, a regulatory region is expected to play a role in prostate cancer progression as well. Thus, in parallel to our MPRA analysis, we also measured the impact of silencing these candidate regions on prostate cancer tumor growth using a loss-of-function strategy in xenograft models (38). We relied on CRISPR-interference, which uses dCas9 fused to a KRAB silencing domain to reduce enhancer activity for the targeted region of interest (39). To systematically target our CDRRs, we engineered an sgRNA library that specifically targets these regions, including non-targeting sgRNA sequences as control (**Fig. 2c**). We then transduced C4-2B CRISPRi-ready cells with this library and compared the representation of guides among the cancer cell populations grown subcutaneously *in vivo,* or grown *in vitro* for a similar number of doublings (**Fig. S2d**). This comparison allowed us to quantify the phenotypic consequences of silencing each of these enhancers. As shown in **Fig. 2d**, there were a number of guides that showed significant association with *in vivo* growth. Moreover, as we had included five independent sgRNAs per regulatory region, we also performed an integrative analysis to combine the phenotypic consequences of guides targeting each region. This allowed us to assign a combined summary phenotypic score to each CDRR. We identified several CDRRs with strong, significant and specific *in vivo* growth phenotypes in the C4-2B prostate cancer cell line (**Fig. S2e**). Similar to our MPRA measurements, this CRISPR-based phenotyping strategy highlighted the enrichment of functional and driver non-coding regions among mCRPC-associated CDRRs.

### Assessing the contribution of individual mutations to CDRR activity

The MPRA and CRISPRi screens described above measured the integrated regulatory and phenotypic impact of hyper-mutated regulatory regions in mCRPC. However, the contributions of individual mutations to the enhancer activity of their containing CDRRs remained unexplored. To shed light on the effects of these mutations at base-resolution scale, we employed two complementary strategies: (i) we used our MPRA assay data to compare the regulatory activity of the reference allele vs. mutant variants, and (ii) we trained a deep learning model to learn the grammar underlying gene expression regulation in prostate cancer. We then used this knowledge to assess the impact of the observed mutations on the expression of its target genes *in silico*.

In the MPRA assay we performed previously, in addition to the reference sequences for each fragment, we had also cloned all mutant variants observed in our patient cohort (**Fig. S3a**). Inclusion of mutant variants enabled us to functionally assess each mutation in CDRRs and measure their phenotypic consequences relative to their reference allele. As shown in **Fig. 3a**, of the more than 350 mutations reliably assayed in the library, about one-third had highly significant impacts on reporter expression relative to the reference allele (FDR <0.01, effect size >1.5-fold). As indicated in **Fig. S3b**, mutations in CDRRs effectively impacted the underlying regions’ activity in prostate cancer cells, highlighting the regulatory consequences of the observed mutations. This observation on its own, however, does not imply that the other two-thirds of mutations are phenotypically neutral. An important caveat here is that our MPRA system removes mutations from their endogenous context and the functionality of some variants may be lost in this transition. Therefore, we took advantage of a machine learning model as a complementary strategy to study these mutations within their larger endogenous context *in silico*.

**Figure 3.**
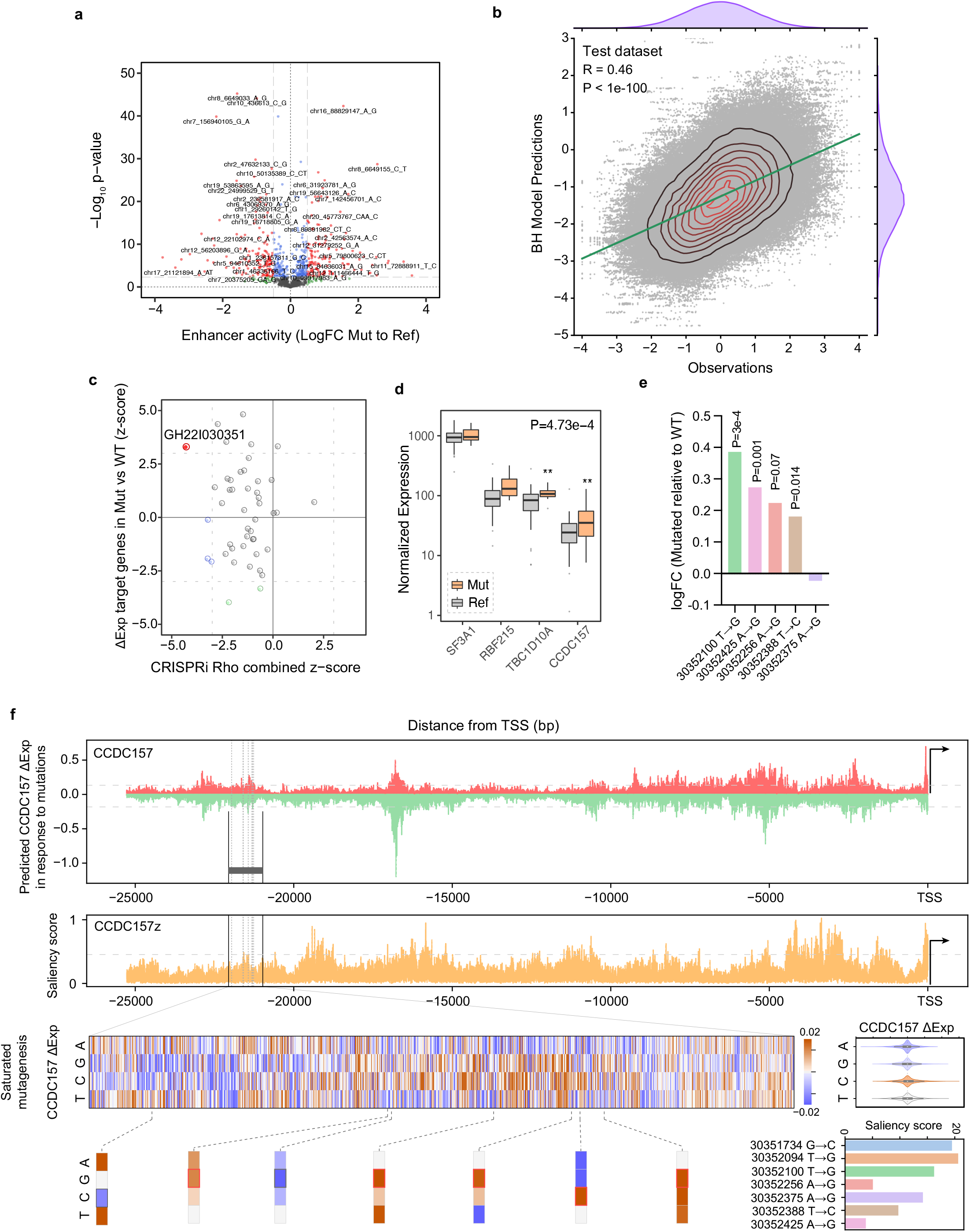
Base-resolution *in vitro* and *in silico* assay reveal the functional consequences of mCRPC-associated mutations. **(a)** A volcano plot demonstrating the impact of individual mutations relative to their reference allele on enhancer activity. **(b)** The overall performance of our Blue Heeler model (BH) in predicting gene expression for held-out instances. **(c)** Comparison of mutational impact on the expression of downstream genes and the overall impact of the mutated regulatory regions based on our *in vivo* screen. A previously annotated enhancer, with genehancer id GH22I030351, shows a strong phenotype in xenografted mice, and patients with mutations in it show generally increased expression in downstream genes. **(d)** Comparing the expression of genes associated with GH22I030351 in mCRPC patient samples with and without mutations in this enhancer. The combined *P*-value shows the overall effect of mutations across all these genes. **(e)** In four out of five cases, measuring the impact of mutations observed in our cohort show a general increase in regulatory activity of GH22I030351 in our MPRA measurements. **(f)** *CCDC157* (ENSG00000187860) promoter sequence, which is immediately downstream of GH22I030351, was used to dissect the impact of mutations *in silico* based on feature attribution scores from our BH model. The top panel shows the results of an *in silico* saturation mutagenesis experiment, in which the impact of every mutation upstream of *CCDC157* on its expression was measured. We observed both gain-of-function and loss-of-function mutations. The regulatory region of interest is shown as a box and the mutations observed in patients are marked by dashed lines. We have also reported saliency scores for this promoter. We further zoomed in on saturation mutagenesis results for our regulatory region of interest to show: (i) the distribution of impact scores for types of mutations, (ii) importance score for loci mutated in patients with the exact mutation shown as a bounded box, and (iii) saliency score associated with each mutated locus.

In recent years, deep-learning-based models have proved successful in linking genotypic variation to phenotypic outcomes. As a result, a number of models have emerged that predict the impacts of single-base substitutions, particularly in non-coding regions, on resulting gene expression (40–44). Here, we developed a base-resolution model that learns the regulatory context of mCRPC in relation to the regulatory activity of promoters/enhancers. This model uses a 2^15^ bp-input promoter sequence on one side and an embedding of the cancer cell state on the other to predict the expression of a given gene (**Fig. S3c**). The promoter sequence (starting ∼32 kb upstream of TSS) is represented as a one-hot encoded 4-channel input, and then processed through a series of convolutional and residual dilation blocks. The resulting sequence embedding is then merged with the output of a cancer state encoder, which provides an embedding of the gene expression profile of each tumor. This cancer-state encoder is pre-trained as a variational autoencoder prior to transfer to the final model. The final layer of the model is a fully-connected layer that predicts expression of a gene given its promoter sequence and the gene expression state of the corresponding sample. The underlying concept is that the convolutional blocks learn the *cis*-regulatory elements and the combinatorial code between them to predict the expression of every gene in a given sample based on the occurrence of these elements along the promoter sequence. Our deep-learning model, which we named Blue Heeler (BH), accomplished this task and predicted gene expression in mCRPC samples using promoter sequences (**Figs. 3b, S3d**). More importantly, it also helped us prioritize functionally relevant mutations and better understand their impact on gene expression control. BH was trained over 87.5% of genes, with the other 10% held out as a test set, and 2.5% for validation (see Methods for more details).

To take a deeper dive and better understand the sequence-function relationships we observed in cells, *in vivo*, and *in silico*, we integrated our results to prioritize the strongest mCRPC-associated regulatory regions. Through this selection process, we nominated a previously annotated enhancer on chromosome 22 as a driver of prostate cancer progression, genehancer ID: GH22I030351 (**Fig. S3e**). Specifically, GH22I030351 showed the most significant enhancer activity after aggregating fragment activity in our MPRA data (**Fig. S2c,** see Methods for aggregation details). Targeting GH22I030351 showed the strongest impact on tumor growth in xenografted C4-2B cells, and mCRPC patients with mutations in this enhancer showed a significant increase in the expression of the genes previously associated with this enhancer (**Figs. 3c-d**). In addition, in almost all cases, observed mutations in this regulatory region significantly increased the activity of this enhancer in our MPRA measurements (**Fig. 3e**). Since this enhancer is ∼20kb upstream of *CCDC157*, we used our pre-trained BH model to analyze this enhancer *in silico*. (We specifically used *CCDC157* from the four gene targets because GH22I030351 strictly falls within the range of distance from the transcription start site that BH is trained on.) First, as expected, we observed that feature attribution scores, as measured by sequence making, sequence variations, and saliency scores, identified GH22I030351 as an important region in regulation of CCDC157 expression (**Fig. 3f**). Moreover, while *in silico* saturation mutagenesis experiments across the *CCDC157* promoter revealed both loss- and gain-of-function mutations, the mCRPC patient mutations in this enhancer were deemed to be largely gain-of-function alterations by the model. This is consistent with our findings from MPRA measurements and the direction of gene expression changes in clinical samples. Together, these observations indicate that GH22I030351 is a strong contender as a non-coding driver in mCRPC by acting as a positive regulator of the expression of its downstream targets.

### SF3A1 and CCDC157 promote prostate cancer downstream of GH22I030351

First, to validate our results from our *in vivo* CRISPR-interference screen, we used a CRISPRi sgRNA construct (the best-performing of the five guides per gene included in our screen) to silence GH22I030351 in C4-2B cells and performed subcutaneous tumor growth assays. As shown in **Fig. 4a**, consistent with the results from our pooled screen, we observed a significant reduction in tumor growth in xenografted mice in GH22I030351-silenced cells. We then asked whether GH22I030351 in fact regulates the expression of its downstream target genes. We silenced GH22I030351 and performed quantitative real-time PCR for the four target genes described for this enhancer, namely *SF3A1*, *CCDC157*, *TBC1D10A*, and *RNF215*. We observed a significant reduction in the expression of SF3A1 and CCDC157, but not TBC1D10A or RNF215 (**Figs. 4b, S4a**). This observation implies that the reduction in tumor growth associated with GH22I030351 resulted from the reduced expression of either, or both, SF3A1 and CCDC157. Interestingly, this observation is consistent with results from whole-genome *in vitro* CRISPRi screens in isogenic LNCaP and C4-2B lines (45). As shown in **Fig. S4b**, sgRNAs that targeted the promoters of *SF3A1* and *CCDC157* resulted in a significant reduction in proliferation in this dataset. However, since these genes share a bidirectional promoter, CRISPRi signals may very well leak from one gene to the other. Therefore, to identify which of these two genes acts as a promoter of prostate cancer growth, we used shRNAs to independently knock down SF3A1 and CCDC157 in C4-2B cells. We used inducible shRNA constructs to measure cancer cell proliferation and colony formation after inducing knockdown of either SF3A1, CCDC157, or both (**Figs. 4c-d**). Interestingly, we observed that constitutive expression of shRNAs against either of these genes was not tolerated by prostate cancer cells, which implies that both of these genes may be acting as drivers. To further confirm this hypothesis, we performed gain-of-function studies by over-expressing these two genes as *trans* genes. As shown in **Fig. 4e**, increasing the expression of either SF3A1 or CCDC157 in C4-2B cells resulted in enhanced tumor growth in xenografted mice. These studies establish GH22I030351 as a major enhancer that simultaneously controls both SF3A1 and CCDC157, which both can act as prostate cancer drivers.

**Figure 4.**
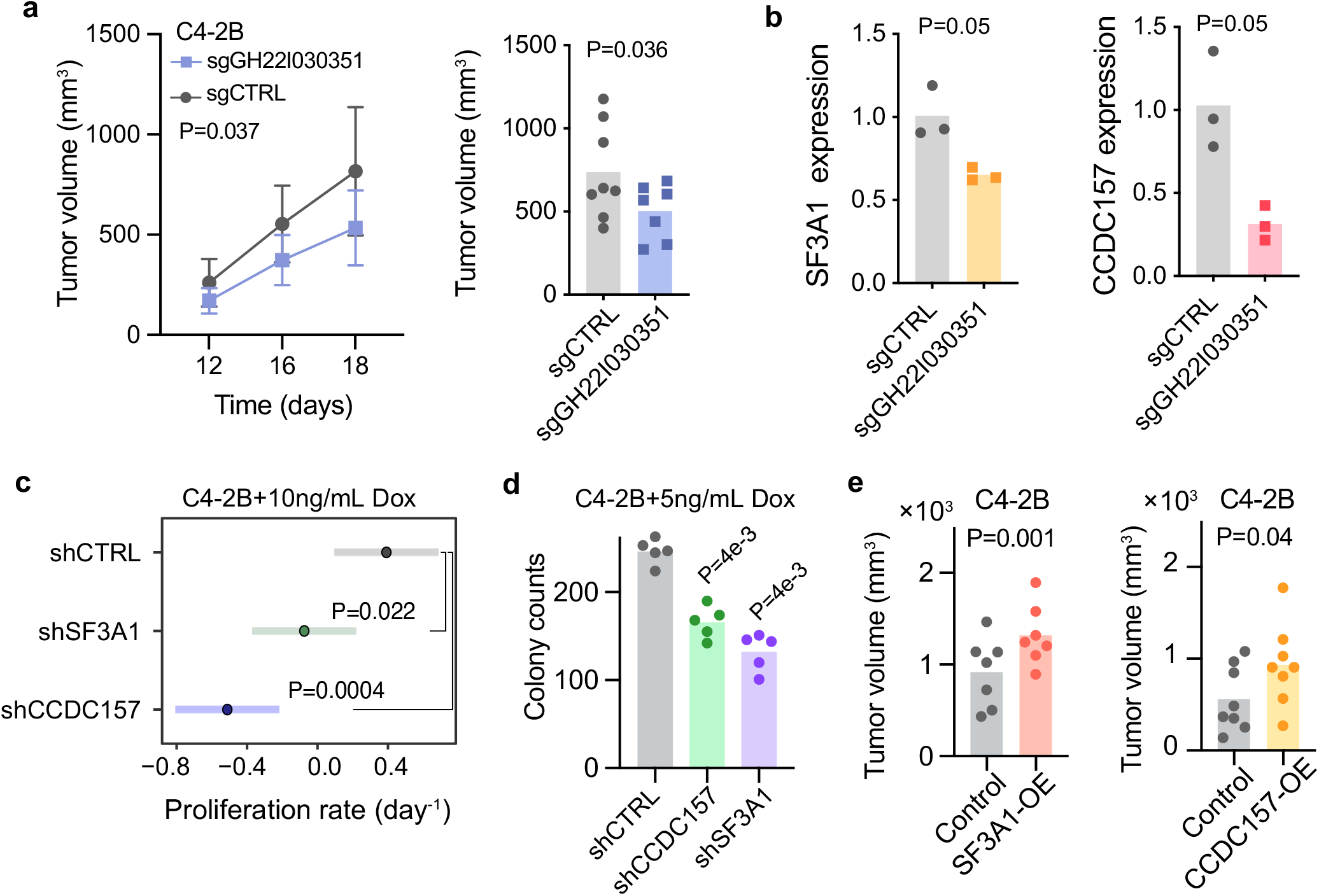
GH22I030351 promotes prostate cancer growth through modulation of SF3A1 and CCDC157 expression. **(a)** Subcutaneous tumor growth in CRISPRi-ready C4-2B cells expressing non-targeting control or sgRNAs targeting GH22I030351. Two-way ANOVA was used to calculate the reported *P*-value. Also shown is the size of extracted tumors at the conclusion of the experiment (day 18 post-injection); *P* calculated using one-tailed *t*-test (n=8 and 7, respectively). **(b)** SF3A1 and CCDC157 mRNA levels, measured using qPCR, in control and GH22I030351-silenced C4-2B cells (n=3). *P* based on a one-tailed Mann-Whitney *U* test. **(c)** Comparison of proliferation rates, as measured by the slope of log-cell count measured over 3 days, for control as well as SF3A1 and CCDC157 knockdown cells (n=6 per shRNA condition). Hairpin RNAs were induced at day 0 and cell viability was measured at days 1, 2, and 3. *P*-values were calculated using least-square models comparing the slope of each knockdown to the control wells. **(d)** Colony formation assay for SF3A1 and CCDC157 knockdown cells in the C4-2B background. Hairpin RNAs were induced at day 0 and colonies counted at day 8. *P*-values were calculated using one-tailed Mann-Whitney U tests. **(e)** Subcutaneous tumor growth in C4-2B cells over-expressing SF3A1 and CCDC157 ORFs in a lentiviral construct. Tumors were measured using calipers at ∼3 weeks post-injection and *P*-values were calculated using a one-tailed Student’s *t*-test.

### SF3A1 over-expression reprograms the splicing landscape of prostate cancer cells

Reprogramming of the alternative splicing landscape is a hallmark of prostate cancer (46). Since SF3A1 is a known splicing factor and a known component of the mature U2 small nuclear ribonucleoprotein particle (snRNP), our observation that SF3A1 up-regulation is implicated in prostate cancer progression further highlights the importance of splicing dysregulations in mCRPC (47-48). We asked whether mutations in GH22I030351, which lead to increased SF3A1 expression, are accompanied by splicing landscape alterations. For this, we used the mixture-of-isoforms (MISO) analytical package (49) to calculate the percent-spliced-in (Ψ) for annotated cassette exons that are expressed in our mCRPC cohort. As shown in **Fig. 5a**, we observed significant alterations in the splicing landscape of cassette exons in GH22I030351-mutated samples, however, this observation on its own does not implicate downstream SF3A1 up-regulation as the immediate cause. While SF3A1 is a canonical component of U2 snRNP, it also directly binds RNA and therefore may influence splicing directly through interactions with target RNAs (50). In order to assess this possibility and draw a more causal link, we decided to specifically focus on transcripts that are directly bound by SF3A1. For this, we used UV cross-linking immunoprecipitation followed by RNA footprinting and sequencing (CLIP-seq) to map SF3A1 binding sites at nucleotide resolution (51). We annotated roughly 40,000 binding sites across the transcriptome, the majority of which fell in intronic regions (**Fig. S5a**). This extensive intronic binding is consistent with the role of SF3A1 as a splicing factor. More importantly, since CLIP-seq provides base-resolution interaction maps, we used high-confidence SF3A1 binding sites to ask whether there were any specific sequence features preferred by SF3A1. As shown in **Fig. 5b**, systematic sequence analysis revealed a significant enrichment of CU-rich elements in SF3A1 sites. Interestingly, it is known that SF3A1 binding to U1 snRNA is directed through an interaction with the terminal CU in the U1-SL4 domain (52). Cassette exons with direct SF3A1 binding also showed increased usage in GH22I030351-mutated tumors (**Figs. 5c, S5b**).

**Figure 5.**
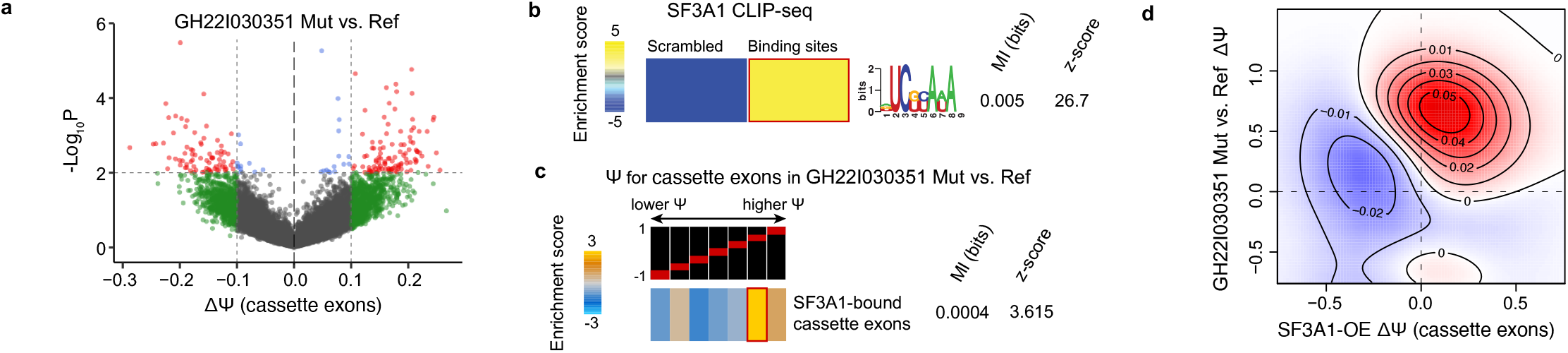
SF3A1 up-regulation results in splicing alterations similar to those observed in GH22I030351-mutated tumors. **(a)** A volcano plot comparing cassette exon usage (percent-spliced-in, Ψ) between tumors with mutations in GH22I030351 relative to other samples in our cohort. Marked are cassette exons with larger than 10% change in Ψ (ranging between -1 to 1) and a *P*-value of <0.01. **(b)** SF3A1 CLIP-seq in C4-2B lines allowed us to identify, at base resolution, high-confidence binding sites of SF3A1 by mapping crosslinking-induced deletions. We used 10nt-long sequences flanking thousands of SF3A1 binding sites to identify sequence preferences for this RBP. To generate a background set of sequences, we also scrambled each binding site while maintaining its dinucleotide content. We used FIRE (37) to discover the most significant sequence motif, and here we report its associated mutual information (MI) and z-score. **(c)** The enrichment of cassette exons bound by SF3A1 among those with higher Ψ in samples with mutations in GH22I030351. For this analysis, we ordered all annotated cassette exons based on their ΔΨ values from -1 (left) to +1 (right). We then grouped them into equally populated bins and assessed the non-random distribution of SF3A1-bound cassette exons across these measurements using MI (31). Individual bins are colored based on their hypergeometric *P*-value as well. **(d)** Comparison of changes in Ψ values in GH22I030351-mutant and SF3A1 over-expression samples. We observed a significant enrichment of SF3A1 binding among those cassette exons that are simultaneously up-regulated in both GH22I030351-mutant and SF3A1 over-expression samples. It should be noted that unbound cassette exons do not show a correlation between these two sets of comparisons.

To confirm this, we performed total RNA sequencing in SF3A1 over-expressing cells, relative to mock-transduced control, in the C4-2B background. As shown in **Fig. S5c**, we observed a number of cassette exons that are significantly up- or down-regulated upon SF3A1 over-expression. More importantly, we observed a significant and clear enrichment of SF3A1-bound cassette exons among those that are up-regulated in SF3A1 over-expressing cells (**Figs. S5d-e**). Finally, comparing the changes in splicing caused by mutations in GH22I030351 to those caused by over-expression of SF3A1 showed that while there was no correlation in alternative splicing patterns across all cassette exons, exons bound by SF3A1 were similarly enriched among the most affected exons in both cases (**Fig. 5d**). Taken together, these observations further highlight a direct link between SF3A1 up-regulation and subsequent RNA binding, and changes in the prostate cancer splicing landscape.

### Putative transcription factors driving GH22I030351-mediated regulation of gene expression

We then sought to identify the upstream transcriptional regulators of SF3A1 and CCDC157 expression that may be impacted by observed mCRPC mutations. We hypothesized that in addition to having a sequence motif match to the GH22I030351 region, given the association of this region with tumor progression, its regulators would also exhibit a metastasis-relevant property, such as increased expression specific to metastatic prostate tumors. While we found 34 transcription factor sequence motifs with significant enrichment at the genomic window intersecting the observed mutations, only 6 were associated with metastatic prostate tumors. We further investigated the top three candidates, SMAD2, TEAD1, and SOX6, and found that the sequence motif match for each of these transcription factors overlapped with mutations observed in our patient cohort (**Figs. 6a**, **S6a**).

**Figure 6.**
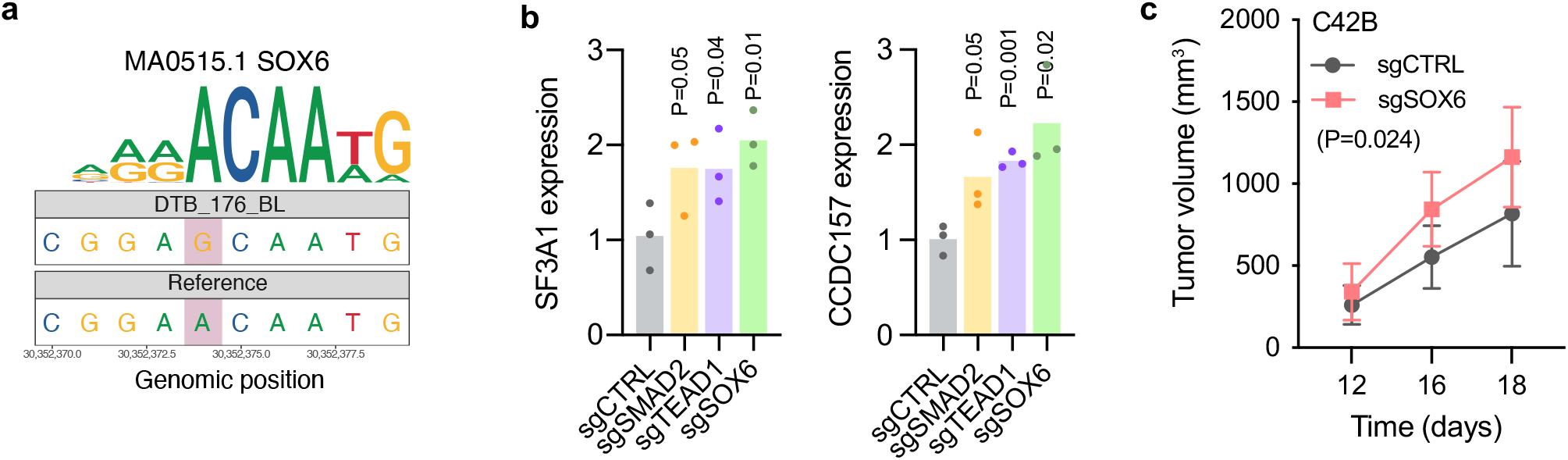
Putative transcription factors that regulate gene expression through GH22I030351. **(a)** A mutation observed in GH22I030351 impacts a SOX6 binding site, likely reducing its binding preference. **(b)** Changes in the expression of SF3A1 and CCDC157 in response to silencing transcription factors we hypothesized to regulate their expression. *P* calculated using a one-tailed Welch’s *t*-test. **(c)** Subcutaneous tumor growth in SOX6 knockdown and control cells in xenografted mice (n=8). *P* calculated using two-way ANOVA using time as a covariate.

To assess the regulatory potential of these transcription factors, we performed CRISPRi-mediated knockdown of each and measured changes in the expression of SF3A1 and CCDC157. For all three transcription factors, SMAD2, TEAD1, and SOX6, we observed a concomitant increase in the expression of these target genes; however, SOX6 silencing showed the strongest effect size for both SF3A1 and CCDC157 (**Fig. 6b**). Consistently, we observed that subcutaneous injection of C4-2B cells with SOX6 knockdown resulted in increased tumor growth in xenografted mice (**Fig. 6c**). In contrast, SMAD2 and TEAD1 knockdown cells did not show a significant change in tumor growth (**Fig. S6b**). We also observed SMAD2 as one of the transcriptional regulators of prostate cancer cells in our MPRA analysis (**Fig. S2b**). Taken together, our observations implicate multiple transcription factors, most notably SOX6, that likely regulate expression of SF3A1 and CCDC157 downstream of GH22I030351.

## Discussion

The oncogenic driver events in non-coding regulatory regions are increasingly gaining recognition with the *TERT* promoter standing out as a prime example (2, 3, 13–15, 40, 53–59). Compared to driver mutations in coding sequences, our understanding of non-coding variants has been hindered by the much larger size of the non-coding genome, the absence of clear direct functional consequences of mutations in non-coding regions, and the limited availability of WGS data. In this study, we have described an integrative computational-experimental framework to systematically identify non-coding drivers of human cancers. This framework combines the power of *in silico* machine learning models with the throughput of massively parallel reporter assays and large-scale *in vivo* genetic screens, and is readily generalizable to other cancer models as well.

Here, we integrated genomic, transcriptomic, epigenomic, and sequence context data via our statistical and deep learning models, MutSpotterCV and DM2D. We used a comprehensive list of covariates and features, and performed a wide range of sensitivity analyses to ensure the robustness of our modeled background mutation rate and computational framework. We then computationally nominated 98 Candidate Driver Regulatory Regions (CDRRs) as potential mutational hotspots in non-coding genomes among >100 mCRPC tumors with downstream gene expression consequences. Next, to functionally validate these CDRRs we used massively parallel reporter assay (MPRA) (30), regulon analysis, and *de novo* motif discovery, and searched for significant associations of CDRRs with enhancer activity. JunD has been demonstrated to play a crucial role in prostate cancer, promoting cell growth (34, 35). Consistent with this, our MPRA results showed that JunD binding is significantly associated with increased enhancer activity. Collectively, we found a strong enrichment of regulatory sequences in CDRRs, providing strong evidence for many CDRRs to play a regulatory role in gene expression in mCRPC. Finally, to evaluate the role of CDRRs in prostate cancer tumor growth, we silenced them using a loss-of-function approach in xenograft models via CRISPRi. This mirrored our MPRA results and revealed a concentration of functional and driver regions in CDRRs associated with mCRPC. We also narrowed our focus to assess the effect of individual mutations at a base-level resolution through a comparative MPRA assay and also our deep learning model, BH, that learns the regulatory context of mCRPC in relation to the regulatory activity of promoter/enhancer sequences.

By integrating our *in silico*, *in vitro*, and *in vivo* results we prioritized the enhancer element GH22I030351 as the most potent mCRPC-associated regulatory region. By silencing GH22I030351, which targets a bidirectional promoter, we were able to uncover SF3A1 and CCDC157 as promoters of prostate cancer progression. Interestingly, genes under the control of bidirectional promoters frequently function as tumor suppressors in various cancer types (60–66). In contrast, we present in this study a gene-pair example that can function as a driver of mCRPC progression. Additionally, previous work suggests that bidirectional promoters are comparatively protected against somatic and epigenetic alterations relative to single promoters (63). Our study shows mutations in proximal non-coding regions as a mechanism of regulating expression of genes under the control of bidirectional promoters.

It is known that change in the splicing landscape is a hallmark of prostate cancer (46), which puts in context our observations of SF3A1 over-expression and its impact on alternative splicing. It was recently shown that the binding of SF3A1 to U1 snRNA is mediated by the CU terminal in the U1-SL4 domain (52). In line with this, we found a significant enrichment of CU-rich elements in SF3A1 sites. Furthermore, cassette exons with direct SF3A1 binding showed increased usage in GH22I030351-mutated tumors. We confirmed this by performing total RNA sequencing in SF3A1 over-expressing cells, relative to control, in the C4-2B background. Interestingly, mutations in another component of the mature U2 snRNP, SF3B1, which have been well-documented in various hematological malignancies, are the most common spliceosomal mutations observed in human cancer (64, 66). These mutations affect coding regions of *SF3B1*, with specific mutations in *SF3B1* associated with well-known clinical outcomes. In addition, *SF3B1* mutations are often gain-of-function, consistent with the tumor growth phenotypes we observed in this study upon over-expressing *SF3A1*. However, coding mutations in *SF3A1* have been noted at comparatively low frequencies across cancer (68). This highlights the strength of our integrated discovery platform for non-coding cancer drivers: though the role of *SF3A1* in cancer progression has remained relatively uncharacterized, our study is the first to show the upstream regulation of *SF3A1* by GH22I030351 and its functional relevance in mCRPC.

Prostate cancer studies previously have identified recurring mutations in pathways involved in androgen signaling and DNA repair, and several genomically distinct classes of prostate cancer involving mutations in transcription factors *SPOP*, *FOXA1*, or *IDH1* have been established (23, 69). Given this diversity in genomic alterations that are present in prostate cancer patient populations, accurate clinical stratification as defined by patient mutational status is essential for proper therapeutic approaches. In this study, we posited that in the absence of a clear functional readout, such as synonymous vs. nonsynonymous mutations in coding sequences, transcriptomic data and comprehensive computational models are essential to understand the pathophysiology of non-coding driver mutations.

During the course of our study, several independent groups have tackled this foundational problem as well. First, a recent pan-cancer study integrated 13 well-established driver discovery algorithms to nominate driver events in coding and non-coding regions in more than 2,600 whole genomes from the Pan-Cancer Analysis of Whole Genomes (PCAWG) dataset across 27 tumor types, including a total of 199 prostate tumors (13, 70). Curiously, their only plausible non-coding driver hit in regulatory regions of prostate tumors was the promoter of the lncRNA gene RP5-997D16.2, having two mutations in their prostate cohort. The authors indicated that they were unable to functionally characterize this non-coding driver, and that there was a lack of overall support for its role based on other evidence. However, by restricting hypothesis testing to boost their statistical power, the authors were also able to find another non-coding hit in the 3’ UTR of the oncogene *FOXA1*. Interestingly, this same region was also tagged as a CDRR in our computational analyses (**Tables S2-3**).

More recently, another pan-cancer study of about 4,000 whole genomes on 19 tumor types (with a total of 341 prostate tumors) from PCAWG and the Hartwig Medical Foundation (HMF) combined statistical significance tests on epigenetic signals, local mutational density, and biological relevance to nominate potential driver events in coding and non-coding regions (3, 71). In this study, the authors primarily used a computational approach and reserved experimental verification only for a single target, namely *XBP1*. In contrast to the higher resolution of our study, the maximum resolution used in that study was a 1-kb tiling window. This study nominated driver events in the coding region, but not the non-coding region, of *SF3B1* in breast, leukemia, and pancreas tumors. Curiously, they also found evidence of strong mutagenic processes, but not driver events, in the vicinity of five prostate-tissue specific genes, namely *ELK4*, *KLK3*, *TMPRSS2*, *ERG*, and *PLPP1*. Of note, we also identified *KLK3* as one of the flanking genes within 15 kb of our non-coding mutational hotspots that we examined for differential gene expression between mutant and reference. However, *KLK3* did not exhibit a significant difference in gene expression levels in our cohort, and thus we excluded this gene from further analysis.

Although these two recent studies had doubled and tripled the whole genome prostate sample sizes compared to ours, their inability to identify any significant driver events in non-coding regions implies that detecting such events in non-coding regions requires a more comprehensive integration of computational and experimental methods. Our results strongly indicate that a computational prioritization fails to paint the full picture, and experimental tools, such as CRISPRi screens and MPRAs, should be part of the discovery platform, rather than a final step for targeted verification of some findings. This goes the other way as well: though previous experimental studies have used experimental approaches like MPRA to study targeted sets of prostate-cancer-relevant cis-regulatory elements (12), we present here a framework for unbiased driver discovery. Moreover, it appears that searching for driver mutations in non-coding genomes may not necessarily derive significant advantage from a pan-cancer cohort. Rather, our study underscores the significance of a targeted cohort with a specific cancer type. Moving forward, we anticipate this integrated framework to be of use for non-coding driver discovery in other cancer patient populations.

## Materials and Methods

### 1. MutSpotterCV

#### 1.1 Data collection and preparation for the MutSpotterCV model

The annotated data for the genomic functional regions were downloaded from three publicly available databases for hg38 as follows. Up/downstream of all genes, untranslated regions, and CpG islands were downloaded from UCSC genome database https://genome.ucsc.edu, with 446,983 entries. Genes were downloaded from ENSEMBL https://www.ensembl.org having a total number of 64,561 entries. Finally, promoters, enhancers, and promoters/enhancers were downloaded from GeneHancer https://www.genecards.org with 250,733 number of entries. These resulted in a total number of 762,277 functional genomic regions which were made consistent in terms of baseness, and subsequently, refined by removing duplicated regions and mitochondrial/unknown chromosomes and random contigs. These regions were further refined by removing very small (<50 bp) and very large (>10,000 bp) regions, resulting in a total of 674,330 annotated functional regions.

There are many overlapping segments among these regions which will bias the downstream analyses, as a mutation can be located in a shared segment and thus counted twice or more, and thus artificially overestimates the mutational density in the region. We thus fragmented overlapping regions using one-hot encoding technique. This technique guarantees that each now-fragmented segment appears only once in the downstream analyses and avoids mutation overcount (**Fig. S1b**). This resulted in 728,208 one-hot encoded, non-overlapping genomic functional regions that are individually labeled by a nine-bit binary digit based on the contribution of each of the nine genomic functional regions (**Fig. S1b**). Each bit would serve as a covariate in the final regression model. Moreover, the length distribution of regions reveals that one-hot encoding produces functional regions with a smoother distribution (**Fig. S1c**).

Next, for each one-hot encoded, non-overlapping functional region we calculated dinucleotide densities and GC content using KENT utility version 403 developed by the UCSC (http://hgdownload.soe.ucsc.edu/admin/exe/linux.x86_64/). We then downloaded 17 available epigenetic features for the cell line PC3 from ENCODE (https://www.encodeproject.org/) with a total number of 18,062,440 entries in the bed format (**Table S1**). These features were then mapped onto our functional regions and subsequently each region was assigned a 17-bit binary number, depending on whether the epigenetic feature existed (1) or not (0) within the region. Each bit represents a covariate in the regression model. Therefore, the total number of covariates in these three classes are 9 + 17 + 17 = 43. However, unsurprisingly, the covariates in sequence context class and GC-content are not independent, nor are the covariates in epigenetic features class. We thus replaced these two classes by their principal components (PCs). As a result, the 16 dinucleotide densities and GC-content were replaced by seven PCs, while 17 epigenetic features were replaced by 10 PCs. In both cases PCs captured >= 80% of variations in data. The selection of PCs encapsulated most of the information embedded in the dinucleotide sequence context. Furthermore, in selecting PCs, we aimed to avoid feature interdependence while simultaneously reducing the number of covariates. This procedure leaves us with a total of 26 new covariates. As can be seen in **Fig. S1d**, all final covariates are statistically significant, meaning they independently contribute to the model prediction.

The small somatic variations, including single nucleotide variations (SNVs) and indels in our cohort are obtained from matched tumor-normal samples as detailed in Quigley et al. (21). Briefly, somatic variations were called by comparing matched normal-tumor samples using Strelka version 2.8.0 (75) and Mutect version 1.1.7 (76), filtered for PASS-designated variations. The total number of small variations in our cohort is 1,890,644 including 1,286,214 SNVs and 604,430 indels. We then cleaned up the somatic variations data by removing mutations on unknown/mitochondrial chromosomes, potential germline mutations (frequency > 1% in the 1000Genome project dataset (77), and single nucleotide polymorphisms recorded in dbSNP (78). This left us with a total number of small variations of 1,874,951 including 1,278,920 SNVs and 596,031 indels. These mutations were mapped onto our one-hot encoded, non-overlapping genomic functional regions using bedtools v2.29.2. Consequently, the mutational density for each functional region was calculated as the number of mutations divided by length to serve as the response variable in our background genomic mutation rate model. Functional regions with zero mutational density were excluded from the rest of the analysis.

#### 1.2 Regression model

With the mutational density as the response variable and 26 covariates, we ran the generalized linear model (GLM), using Gamma distribution for the error structure with the default inverse link function and a variance proportional to μ^2^(with μ being the expected value of the response) in R version 4.0.0. We used a power transformation of the response variable (mutational density) to ensure that the residuals followed a Gamma distribution, and subsequently verified that Gamma was the closest known distribution to our empirical data via a Cullen-Frey graph using the package fitdistrplus version 1.1-1 in R.

#### 1.3 Statistical outlier detection

By systematically comparing the observed vs. expected mutational density, one can determine statistical outliers which serve as the first set of initial candidates for mutation hotspots in this work. Our criteria for a region to be a statistical outlier were i) to harbor at least three mutations ii) the deviance residual of the mutational density be at least one interquartile above the upper quartile (79). These criteria marked 1,780 functional regions as statistical outliers (**Fig. 1b**) which served as the initial set of candidates of being mutational hotspots within the non-coding regulatory regions. Due to the exploratory nature of our analysis, we relaxed multiple testing corrections for outlier detection.

#### 1.4 Copy number alterations

We quantified the sensitivity of the MutSpotterCV’s predictions to the copy number alteration, as this feature is widely present in our cohort (21). We performed this by adding copy number alterations as continuous predictors to the regression model. To do so, we took the DNA copy number variants that had been computed in our cohort binned into windows of 3Mbp by using Canvas version 1.28.0-O01073 (80) and Copycat (https://github.com/chrisamiller/copyCat). First, we mapped the binned windows into our functional regions, and then for each region we replaced the copy number variants by five quantiles, i.e., min, 1^st^ quartile, median, 3^rd^ quartile, and max. This procedure adds five predictors to the original regression model. Nevertheless, there was no significant change in the final predicted statistical outliers in the presence of copy number variations as extra predictors (**Figs. S1e-f**).

#### 1.5 MutSpotterCV on coding sequence

Additionally, to benchmark the MutSpotterCV, we evaluated the model on the coding sequences in our cohort, to ensure that we can recapitulate known recurrently mutated genes as mutational hotspots. In our earlier study we had identified 72 oncogenes in mCRPC (21). Mutational hotspots predicted by the MutSpotterCV in the coding regions capture 64% of previously identified oncogenes in mCRPC (*p* < 10^-16^, hypergeometric test). The absence of a perfect overlap is expected as MutSpotterCV does not currently incorporate structural variations.

#### 1.6 Integration with gene expression data

To find the association of statistical outliers with gene expression in our cohort we first find genes in the 15k bp flanking regions of either ends of all regions. There are a total of 1,692 genes in the flanking regions of 1,264 non-coding mutational hotspots. Notably, not every non-coding mutational hotspot is proximal to a gene. For every statistical outlier, we grouped the cohort into mutation-free (reference) and mutation-bearing (mutant) patients, i.e. patients who do not, or do, have mutations in that non-coding mutational hotspot. Subsequently, for every flanking gene we performed differential gene expression analysis using DESeq2 version 1.28.1 (29).

We find 160 genes with significant change in their expression levels in two groups of patients (*P* < 0.05) proximal to 152 non-coding mutational hotspots. We did not perform multiple testing correction, as, on average, there is rarely more than one gene located in the vicinity of each non-coding hotspot. We then further refined the list by setting the minimum number of mutated patients per region to four, which resulted in 104 flanking genes in the vicinity of 98 non-coding regulatory regions, termed candidate driver regulatory regions (CDRRs), which harbor a total of 885 mutations (**Tables S3-4**). Tumor purity was not a major concern in our analyses as samples were isolated using laser capture microdissection (18).

#### 1.7 Model robustness

We evaluated how the model performance and final results are affected by using different epigenetic data sets from two different cell lines, LNCaP and A549 (ENCODE database). As indicated in **Figs. S1g-j,** we found that using these data sets did not significantly change the model’s performance or the identified mutational hotspots compared to using the default cell line (PC3), indicating collinearity and redundancy of genomic measurements.

### 2. DM2D and the Blue Heeler model

DM2D is a deep convolutional neural network to predict the mutational density values as a function of the underlying DNA sequence and broad functional sequence categories, namely “Gene”, “Enhancer”,”downstream” and “upstream” of genes, “UTR”, “Promoter”, “CpG” island, and “PromHancer” (promoter or enhancer). The “sequence encoder”, with a 7-channel input (3 epigenetic signals and 4 one-hot encoded sequence) contained four convolution blocks, with (16, 32, 32, 32) filters and (4,25,25,25) kernel sizes. All blocks applied batch normalization, rectified linear units, max pooling (window sizes of 4, 10, 10, 10), and 0.25 dropout. The resulting tensors were flattened, concatenated to a one-hot encoded sequence category (size 9), and passed on to fully connected layers with size 24, 12, and 1 respectively. All layers applied batch normalization, rectified linear units, and dropout (0.1). The final layer predicted the mutational density values. For training a Nadam optimizer was used with learning_rate= 0.001, clip_norm=0.5, and clip_value=1. We used MSE as the loss function and trained the model for 20 epochs with a batch size of 128. 15% of samples were held-out as a validation set.

Our Blue Heeler (BH) model is inspired by Basenji (41), with multiple convolutional and dilated convolutional layers. BH contains two inputs, a one-hot encoded sequence input and a sample state input. The former is passed a 2^15^ kb long sequence and the latter a 256-dimensional embedding. For each sample, this embedding was generated using a variational autoencoder with a hidden layer of size 2560, and applying batch normalization and rectified linear units (except for the final layer in the decoder). Expression values were pre-processed by applying rank-based inverse normal transformation prior to training. The Pearson correlation between the reconstructed gene expression values across >100 samples and their input values was on average 0.92. <u>Augmentation:</u> the training data loader, which iterates through promoter sequences of genes, randomly selects one of the samples and uses its embedding as input to the sample state module. Similarly, the promoter sequence, or its reverse complement (with a 50:50 chance) is transformed to a one-hot encoded tensor that is passed on the sequence encoder. <u>Task:</u> the model is then trained to predict the expression of the input gene in the context of the randomly selected sample. <u>Convolutional blocks:</u> four convolutional blocks with (64, 32, 32, 32) filters and (16,8,8,8) kernel sizes. All blocks applied batch normalization, Gaussian error linear units, 0.2 dropout, and max pooling of (16,8,8,8). <u>Dilated</u> <u>convolutional blocks:</u> four densely connected dilated layers with 32 filters and kernel size of 3 and dilations of 2^j^ (where j is the dilated layer number) to increase the receptive field of the sequence encoder. These layers also apply GELU and batch normalization. <u>Regression head:</u> fully connected layers with 1056 and 64 hidden sizes were used to connect the output of the sequence and sample state encoders to the regression head. <u>Training</u>: an Adam optimizer with learning_rate=0.001 and clip_grad_norm of 10 was used to minimize an MSE loss. The model was trained for 60 epochs; 10% of genes were held out as a test set, and 2.5% for validation. The remainder were used for training. The performance of the model was assessed using Pearson correlation applied to all the held-out genes across all samples.

### 3. Sequence motif analysis

We used FIMO (v5.3.2) (81) and JASPAR database core vertebrate non-redundant set of motifs (82)to identify all of the sequence motif matches at the genomic window chr22:30351638–30352714 (hg38 assembly) overlapping the 9 single nucleotide polymorphisms.

We performed DESeq2 (v1.28.1) differential gene expression analysis comparing metastatic to the primary tumors and found that 6 of the 34 transcription factors which have a sequence motif match to the enhancer are significantly upregulated in the metastatic tumors. These included SOX6, SMAD2, TEAD1, PBX3, TEAD2, and SMAD3. We chose the top 3 (SOX6, SMAD2, and TEAD1) for *in vitro* validation.

### 4. Library cloning and sequencing validation

For our MPRA library, a library consisting of 3665 oligos targeting nominated regions was designed and ordered from Twist Biosciences. The pool was resuspended to 5ng/μL final concentration in Tris-HCl 10mM pH 8, and a qPCR to determine Ct to be used for downstream library amplification was performed (forward primer: ATTTTGCCCCTGGTTCTTCCAC, reverse primer: CCCTAAGAAATGAACTGGCAGC) using a 16-fold library dilution.

The library was then amplified via PCR, and ran out on a 2% agarose gel to check library size (expected band of 84bp). PCR product was then cleaned up using a DNA Clean and Concentrator kit-5 (Zymo Research Cat. #D4003), and eluted in 15μL H_2_O. Cleaned product was digested overnight using FD Bpu1102I (Thermo Fisher Cat. #FD0094), and then further digested for 1hr using FD BstXI (Thermo Fisher Cat. #FD1024). Inserts were then ligated into pCRISPRi/a v2 backbone in a 50ng reaction with 1:1 insert:backbone ratio for 16hrs 16C. Ligated products were then ethanol-precipitated overnight at -20C, cleaned, and then transformed into 100μL NEB Stables (NEB Cat. #C3040H), followed by maxiprep plasmid isolation.

For sequencing validation, 1μg plasmid DNA was then digested in 50μL volume for 1hr with FD BstXI (Thermo Fisher Cat. #FD1024). Digested plasmid DNA was then Klenow-extended using added UMI linker (sequence: CTCTTTCCCTACACGACGCTCTTCCGATCTNNNNNNcttg), and then cleaned up using a Zymo DNA Clean & Concentrator-25 kit (Zymo Research Cat. #D4033). Indexing PCR (forward primer: AATGATACGGCGACCACCGAGATCTacactctttccctacacgacgctc; reverse primer: CAAGCAGAAGACGGCATACGAGATGATCTGGTGACTGGAGTTCAGACGTGTGCTCTTCCGATcgactcggtgccactttttc) was then performed in 30μL final volume, followed by gel purification (Takara Bio Cat. #740609.50). Samples were then pooled and sequenced on a lane of HiSeq 4000 SE50 at the UCSF Center for Advanced Technology (CAT).

### 5. Viral transductions

3 million HEK293Ts were seeded in a 15cm plate. 24hrs later, HEK293Ts were transfected with TransIT-Lenti (Mirus Bio Cat. #Mir6603) reagent. Viral supernatant was harvested, aliquoted, flash-frozen, and then stored -80C for long-term storage.

100K C4-2B CRISPRi cells were then seeded in a 6-well plate for viral titering. Using a range of 100-, 200-, and 400μL viral supernatant, cells were transduced, adding polybrene to 8ug/mL final concentration. 48hrs post-transduction, cells were passed through flow cytometry on the FACS Aria II in the UCSF CAT, and %BFP+ was recorded.

### 6. Cell preparation for subcutaneous injection

6 million C4-2B CRISPRi cells were seeded into a 15cm plate and allowed to grow overnight. On the following day, 5.55mL of lentivirus was added to cells (target 33% MOI), with polybrene added to final concentration 8ug/mL. Media was then changed 24hrs post-transduction, and puromycin was added 72 hrs post-transduction to final concentration 2ug/mL. Transduced C4-2B CRISPRi cells were then partitioned into 3 arms for an in vivo CRISPRi screen. 200K cells were split into a 15cm plate for *in vitro* long-term passage, for purposes of growth normalization. 200K cells were pelleted and frozen at -80C for downstream gDNA extraction. 9 million cells were resuspended to final concentration 1 million cells/50μL in 1:1 PBS/matrigel. Bilateral subcutaneous injections in 50μL final volume were then performed in male, 8-12 week-old age-matched male NOD *scid* gamma (NSG) mice (n = 3).

### 7. Tumor gDNA extraction and library preparation

Tumors were then harvested 4 weeks post-injection and processed using Quick-DNA midiprep plus kit (Zymo Research Cat. #D4075). For each processed tumor, genomic DNA was digested in 15ug-scale, 50μL volume reactions with FD BstXI. Digested genomic DNA was then Klenow-extended using added UMI linker (sequence: CTCTTTCCCTACACGACGCTCTTCCGATCTNNNNNNcttg), and then cleaned up using a Zymo DNA Clean & Concentrator-25 kit (Zymo Research Cat. #D4033). Indexing PCRs (forward primer: AATGATACGGCGACCACCGAGATCTacactctttccctacacgacgctc; reverse primer: CAAGCAGAAGACGGCATACGAGATGATCTGGTGACTGGAGTTCAGACGTGTGCTCTTCCGATcgactcggtgccactttttc) were then performed in 30μL final volume, followed by gel purification (Takara Bio Cat. #740609.50). Samples were then pooled and sequenced on a lane of HiSeq 4000 SE50 at the UCSF Center for Advanced Technology (CAT).

### 8. LentiMPRA library cloning

LentiMPRA was performed according to Gordon et al. Briefly, a CRS library was designed and ordered through Twist Biosciences. A first-round PCR reaction was performed to add vector overhang sequence upstream and minimal promoter and adaptor sequences downstream of the CRSs. PCR products were then combined, and cleaned up using 1:1 HighPrep PCR reagent (MagBio Genomics Cat. #AC-60050), eluting in 50μL elution buffer. A second round of PCR was then performed to add a 15-bp barcode and vector overhang sequence downstream of the first-round PCR fragment. PCR products were then combined and ran on two 1.5% TAE-agarose gels, and the resulting band at 419 bp was gel excised and purified using the QIAquick Gel Extraction Kit (Qiagen Cat. #28706X4), eluting in 50μL elution buffer. Resulting DNA was purified using 1.2:1 HighPrep PCR reagent. pLS-SceI backbone was then digested with AgeI-HF (NEB Cat. # R3552S) and SbfI (0.5U/uL) overnight, and then purified using 0.65:1 HighPrep PCR reagent. Linearized pLS-SceI and insert DNA was then recombined using NEBuilder HiFi DNA Assembly Master Mix for 60 min at 50C, and resulting product purified using 0.65:1 HighPrep PCR reagent. Undigested vector was then cut using I-SceI for 1 hr, and resulting DNA purified using 1.8:1 HighPrep PCR reagent, eluting in 20μL elution buffer.

For electroporation, 100ng of recombination product was then added to 100μL of NEB 10-beta electrocompetent cells. Electroporation was conducted in a Gemini X2 electroporator and cells were shocked with 2.0kV voltage; 200 ohms resistance; 25 uF capacitance; 1 pulse; 1 mm gap width. Cells were then grown in 1mL fresh Stable Outgrowth Medium for 1 hour 37C with agitation, and 2μL of bacteria were diluted in 400μL LB medium + 100 mg/mL carbenicillin for colony counting. Undiluted bacteria were plated onto other carbenicillin plates and grown at 37C overnight. 8 colonies were chosen from the dilution plate and sent for Sanger sequencing. 5mL LB media was added to each plate for scraping using a cell lifter, and plasmid was purified using the Qiagen Plasmid Plus Midi Kit.

### 9. LentiMPRA CRS-barcode association sequencing

PCR to add P5 flow cell sequence and the sample index sequence upstream and P7 flow cell sequence downstream of the CRS-barcode fragment was performed using primers pLSmP-ass-i741 and pLSmP-ass-gfp. PCR products were then combined and gel extracted (470bp) under blue light, followed by purification using QIAquick Gel Extraction Kit. DNA was then purified using 1.8X HighPrep PCR reagent, and DNA was sequenced using a MiSeq v2 (15 million reads) kit using custom primers pLSmP-ass-seq-R1 (CRS upstream forward), pLSmP-ass-seq-R2 (CRS downstream reverse), pLSmP-ass-seq-ind1 (Barcode forward), and pLSmP-rand-ind2 (sample index) as described previously.

### 10. Lentivirus packaging

10 million 293T cells were seeded into a 15-cm plate and incubated for 2d. Transfection was then carried out as described previously, using 60μL EndoFectin, 10 μg plasmid library, 6.5 μg psPAX2, and 3.5 μg pMD2.G. Cells were incubated for 14 hours and then media was replaced with 20mL DMEM supplemented with 40μL ViralBoost reagent, and incubated for 48 hours. GFP expression was confirmed using fluorescence microscopy and viral supernatant was then filtered using a 0.45uM filter. Supernatant was concentrated using 1⁄3 volume Lenti-X concentrator reagent, centrifuging for 1500g 45 mins 4C and resuspending the resulting lentivirus pellet in 600μL DPBS.

### 11. Lentivirus titration

100K C4-2B cells were seeded into wells of a 6-well plate. To calculate viral titer, lentiviral library was then infected in a 2-fold upwards range (0, 1, 2, 4, 8, 16, 32, 64uL), gDNA was extracted, and qPCR was performed to determine MOI for each lentiviral library condition.

### 12. Lentivirus infection and library preparation

Using a target of 100 integrations per barcode, 1.1 million C4-2B cells were seeded in a 10cm plate, in three biological replicates. Cells were incubated overnight and culture media was refreshed with polybrene at 8 ug/mL final concentration. 87μL virus was then added to plates and culture media was refreshed with no polybrene the following day. GFP fluorescence was confirmed 2d after, and culture media was removed. Cells were washed 3 times with DPBS and the AllPrep DNA/RNA Mini Kit was used to simultaneously extract DNA/RNA from plates, eluting DNA/RNA fractions in 30μL Buffer EB/RNAase-free H20 respectively. RNA samples were then treated with DNAse and reverse-transcription (RT) reactions were performed in 8-strip PCR tubes. These reactions add a 16-bp UI and P7 flowcells sequence downstream of the barcode, using low-complexity amounts as previously described.

DNA samples were then diluted to 120ng/μL final concentration. 100μL of DNA or RT products respectively (for 12 μg DNA or entire RT product) were then used for a first-round PCR reaction to add the P5 flow cell sequence and sample index sequence upstream and a 16-bp UMI and P7 flow cell sequence downstream of the barcode. DNA was then purified using 1.8X HighPrep PCR reagent and eluted in 60μL elution buffer. A preliminary qPCR reaction was set up to find the number of PCR cycles required for the subsequent second-round PCR reaction with P7 and P5 primers. 23 cycles were then used for the second-round PCR reaction, DNA was purified in a 1.8X HighPrep PCR reagent clean-up, and sample run on 1.8% wt/vol agarose gel. The band at 162 bp was excised and purified using the QIAquick Gel Extraction Kit and purified 1.8X. DNA and RNA samples were then pooled in a single LoBind tube with 1:3 ratio, and final sequencing library sent out to the Center for Advanced Technology (CAT) at UCSF for sequencing on two HiSeq 4000 lanes.

### 13. CLIP-seq of SF3A1 in C4-2B CRISPRi cells

#### UV-Crosslinking

Six 15cm plates of C4-2B CRISPRi cells were seeded for a total of 3 biological replicates. Cells were then harvested 48 hours later and then were crosslinked on a 254nm UV crosslinker set to 400mJ/cm^2^, transferred to 15mL tubes, spun at 1500xg 4C for 2 mins, and then frozen as dry pellets at -80C for long term storage.

#### Bead Preparation

For bead preparation, 60μL Protein A beads were then washed in 2X_[H1]_ in low salt wash buffer (1X PBS, 0.1% SDS, 0.5% sodium deoxycholate, 0.5% IGEPAL CA-630), adding 2μg anti-SF3A1 (Proteintech Cat. #15858-1-AP) and then rotating at 4C for 1hr. For cell lysis, cells were then resuspended in 600μL cold low salt wash buffer + 6μL SuperaseIN (Invitrogen Cat. #AM2696) + 1X protease inhibitor cocktail (Thermo Sci Cat. #78425) and incubated on ice for 10 mins.

#### RNase Treatment and Immunoprecipitation

Cells were then equally divided and treated with either 20μL RNase high mixture (RNase A 1:3,000 + RNaseI 1:10) or 20μL low mixture (RNase A 1:15,000 + RNaseI 1:500) and incubated at 37C for 5 mins, and then combined and spun at 4C max speed for 20 mins. Clarified supernatant was added to prepared beads and rotated end-over-end at 4C, for 2 hours. Beads were collected on magnet and washed 2X with 1mL cold low salt wash buffer, 2X with 1mL high salt wash buffer (5X PBS, 0.1% SDS, 0.5% sodium deoxycholate, 0.5% IGEPAL CA-630), and then 2X with 1mL cold PNK buffer (50 mM Tris-HCl pH 7.5, 10 mM MgCl_2_).

#### RNA Dephosphorylation

For RNA dephosphorylation, 2.5μL 10X PNK buffer (500mM Tris pH6.8, 50mM MgCl2, 50mM DTT), 2μL 10X T4 PNK (NEB Cat. #M0201L), 0.5μL SuperaseIN, 20μL nuclease free water was added per reaction, and incubated at 37C for 20 mins in a thermomixer (mix 1350 rpm 15s/5 mins rest). Beads were then washed 1X with 1mL PNK buffer, 1X with 1mL high salt wash buffer, and 2X with 1mL PNK buffer.

#### PolyA-tailing, N3-dUTP end labeling, and Dye labeling

RNP complexes were then polyA-tailed by addition of 0.8μL yeast PAP (Jena 600U/ul), 4μL 5X yeast PAP buffer, 1μL 10 mM ATP (unlabeled), 0.5μL SuperaseIN, 13.7μL nuclease free water, and incubated at 22C for 5 mins in thermomixer (shake 1X 15s 1350 rpm). After 5 mins incubation, beads were washed 2X with 500μL cold high salt buffer, then 2X 500μL cold PNK buffer. Samples were then N3-dUTP labeled with 0.4μL yeast PAP, 2μL 5X yeast PAP buffer, 0.25μL SuperaseIN, 2μL 10mM N3-dUTP, 5.35μL nuclease free water, and incubated for 20 mins at 37C in a thermomixer with intermittent shaking (15s/5 mins rest, 1350 rpm). Samples were then washed with 2X 500μL cold high salt wash buffer, then 2X with 200μL cold 1X PBS. For dye labeling of N3-labeled RNA, 20μL 1mM 800CW DBCO in PBS was then added, and incubated in a thermomixer protected from light at 22C for 30 mins with intermittent shaking (15s/5 mins rest, 1350 rpm). Beads were then washed 1X with 500μL high salt wash buffer, then 1X with 500μL PNK buffer and then resuspended in 20μL loading buffer (1X NuPAGE loading buffer + 50 mM DTT diluted in PNK buffer), and then heated at 75C for 10 mins shaking at 1000rpm, protected from light. Supernatant were transferred to clean microfuge tubes.

#### PAGE and transfer

Samples were then run on a 12-well Novex NuPAGE 4-12% Bis-Tris gel (1mm thick) at 180V for 90 mins along with IR-labeled protein standard in 1X MOPS running buffer at 4C, light-protected. Gel was then transferred to protran BA-85 nitrocellulose membrane in Novex X-cell apparatus using 1X NuPAGE transfer buffer with 15% EtOH for 75 mins at 30V. Membrane was then rinsed in PBS, and imaged with a Licor Odyssey instrument.

#### Proteinase K digest and RNA capture

Nitrocellulose membrane was excised at the expected range size (140-150kDa) for SF3A1, to capture RNA-protein complexes. Membrane was placed into a clean microfuge tube, and 200μL Proteinase K digestion buffer (100mM Tris-HCl pH 7.5, 100nM NaCl, 1mM EDTA, 0.2% SDS), 12.5μL Proteinase K, was added. Samples were then incubated at 55C for 45 mins in a thermomixer at 1100rpm. Samples were spun and the supernatant was transferred to clean microfuge tubes, and the final solution was adjusted to ∼500mM NaCl by adding 19μL 5M NaCl and 11μL nuclease free water. Salt-adjusted solution was then added to pre-washed oligo-dT dynabeads, incubating for 20 mins at 25C in a thermomixer with intermittent shaking (1350rpm, 10s/10 mins, 300 rpm remainder of time). Beads were then washed 2X with 100μL cold high salt wash buffer, 2X with 100μL cold PBS. Samples were eluted from beads with 8μL of TE buffer (20 mM Tris-HCl pH 7.5, 1mM EDTA), heated at 50C for 5 mins, and 7.5μL of supernatant was transferred into a clean PCR tube on ice.

### 14. cDNA synthesis and PCR

For annealing, to 7.5μL eluted RNA 2.5μL smRNA mix 1 (Takara Cat. #635031) and 1μL 10uM UMI RT primer (seq: CAAGCAGAAGACGGCATACGAGATNNNNNNNNGTGACTGGAGTTCAGACGTGTGCTCTTCCGATCTTTTTTTTTTTTT TT) were added, heated at 72C 3 mins in a thermocycler, and then placed on ice for 5 mins. 9μL RT mix (6.5μL smRNA Mix 2, 0.5μL RNAse inhibitor ((Invitrogen Cat. #AM2696), 2μL PrimeScript RT (200U/ul)) was then added to samples on ice, and the following program was run: 42C 60 mins, 70C 10mins, 4C hold.

For indexing PCR, 78μL PCR mix (24μL H20, 50μL 2X SeqAmp CB PCR buffer (Takara Cat. #638526), 2μL SeqAmp DNA polymerase ((Takara Cat. #638509), 2μL 10uM universal reverse primer (seq: CAAGCAGAAGACGGCATACGAG)) was added to each cDNA sample, followed by 2μL of 10uM indexed forward primer (seq: AATGATACGGCGACCACC). The following program was run for: 98C 1 min, [98C 10s, 60C 5s, 68C 10s, repeat 18X], 4C hold. Product was size-selected using 1.1X bead: PCR ratio, and final product was eluted in 16μL H20. Samples were quantified via Agilent Tapestation 4200 and submitted for sequencing on a lane of HiSeq 4000 SE 50.

### 15. Cell growth assays

For assaying cell proliferation, CellTiter-Glo 2.0 Cell Viability Assay (Promega Cat. #G9241) was used. 1K C4-2B cells were seeded per well in 3 separate opaque 96-well plates for luminescence measurement at days 1, 2, and 3. 6 wells were seeded per cell condition in 100μL volume media. 24h after seeding, media was replaced with fresh media containing doxycycline at 10 ng/mL final concentration. Cells were then harvested according to manufacturer’s protocol. Briefly, CellTiter-Glo 2.0 Reagent and cell plates were equilibrated to RT 30 mins prior to use. 100μL CellTiter-Glo 2.0 Reagent was then added via multichannel to each well and mixed at 300 rpm for 2 mins at RT; the plate was incubated for 10 minutes at RT, covered. Plate luminescence was then recorded on a SpectraMax iD5 multiplate reader.

For colony formation assay, 2.5K C4-2B cells were seeded in triplicate in a 6-well plate. 24h after seeding, media was replaced with media containing doxycycline at 5 ng/mL final concentration. 8 days after doxycycline induction, colonies were stained and imaged. Briefly, media was removed and cells were washed with 1mL PBS at RT. Cells were then fixed in 4% PFA (Alfa Aesar Cat. #43368-9L) for 10 minutes at RT, and then stained in 0.1% crystal violet (Sigma-Aldrich Cat. #V5265-250ML) for 1h at RT. Wells were then washed 3X with ddH20 at RT until colonies were visible. Colonies were imaged on an Azure c200 and counted.

### 16. Cell culture

All cells were cultured in a 37°C 5% CO2 humidified incubator. The C4-2B prostate cancer cell line was cultured in RPMI-1640 medium supplemented with 10% FBS, glucose (2 g/L), L-glutamine (2 mM), 25 mM HEPES, penicillin (100 units/mL), streptomycin (100 μg/mL) and amphotericin B (1 μg/mL) (Gibco). All cell lines were routinely screened for mycoplasma with a PCR-based assay.

### 17. RNA-seq

RNA-seq was done on SF3A1 over-expression and control cell lines. RNA was extracted from samples by column clean up (Zymo Quick-RNA Microprep Kit). RNA-seq libraries were prepared from these samples using the SMARTer® Stranded Total RNA-Seq Kit v3 - Pico Input Mammalian (Takara) kit according to manufacturer’s instructions. Sequencing was performed on an Illumina NextSeq 5000.

## List of Supplementary Materials

Figs. S1-S6

Tables S1-S4

## Acknowledgements

We thank members from the Ahituv lab for assisting BJW with lentiMPRA cloning, specifically Gracie Gordon and Dianne Laboy Cintrón. We thank the Laboratory Animal Resource Center (LARC) at UCSF.

## Data and materials availability

MutSpotterCV, DM2D, and BlueHeeler are available at github.com/goodarzilab. RNA-seq, MPRA, and CRISPR screening data generated as part of this study is deposited to Gene Expression Omnibus, and is available under the accession ID GSEXXXX.

