## Supplementary material for "Integrative identification of non-coding regulatory regions driving metastatic prostate cancer": Figs. S1-6

### Supplementary Figures

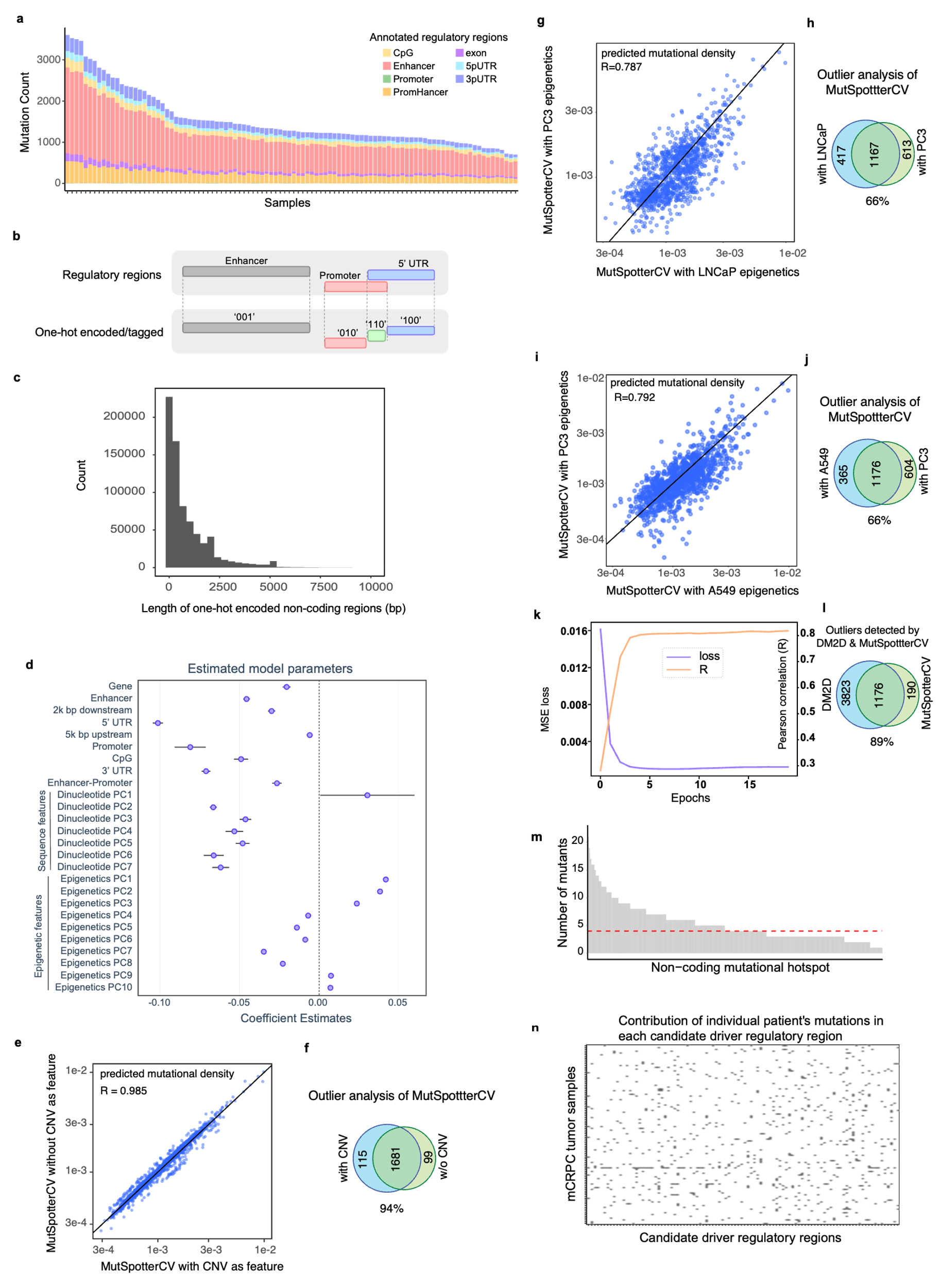

##### Figure S1. MutSpotterCV effectively predicts the background mutational density in regulatory regions.

**(a)** Number of mutations (including SNVs and indels) per sample per regulatory region sorted in a descending order. As shown here, the total number of mutations were largely similar across these patients; however, one hypermutated sample was removed to avoid bias. Each color shows the proportion of mutation counts in the corresponding regulatory region. **(b)** Depiction of our one-hot encoding strategy to uniquely tag overlapping functional genomic regions. **(c)** Distribution of the lengths of regulatory regions after one-hot encoding as inputs for the MutSpotterCV. **(d)** The forest plot of the final MutSpotterCV model covariates showing all features are significant in the final prediction. **(e-f)** Comparison of MutSpotterCV with and without copy number (CN) as an additional model parameter. Shown are (e) the correlation between predicted mutational densities in outliers, with and without CN, and (f) the Venn diagram of the MutSpotterCV-predicted outliers with and without CN. **(g-j)** Comparison of MutSpotterCV using epigenetic features from PC3 cell lines (default) vs. LNCaP or A549. Shown are the correlation between predicted mutational densities in outliers when using PC3 vs. (g) LNCaP or (i) A549. The corresponding Venn diagrams of the outliers are shown in (h) and (j). **(k)** The validation loss and Pearson correlation for the DM2D model as a function of epochs. **(l)** Venn diagram of non-coding mutational hotspots independently detected by MutSpotterCV and DM2D. **(m)** Number of mutants per non-coding mutational hotspots. We required each non-coding hotspot to include at least four mutants (dashed red line) **(n)** Distribution of mutations in CDRRs in patient samples.

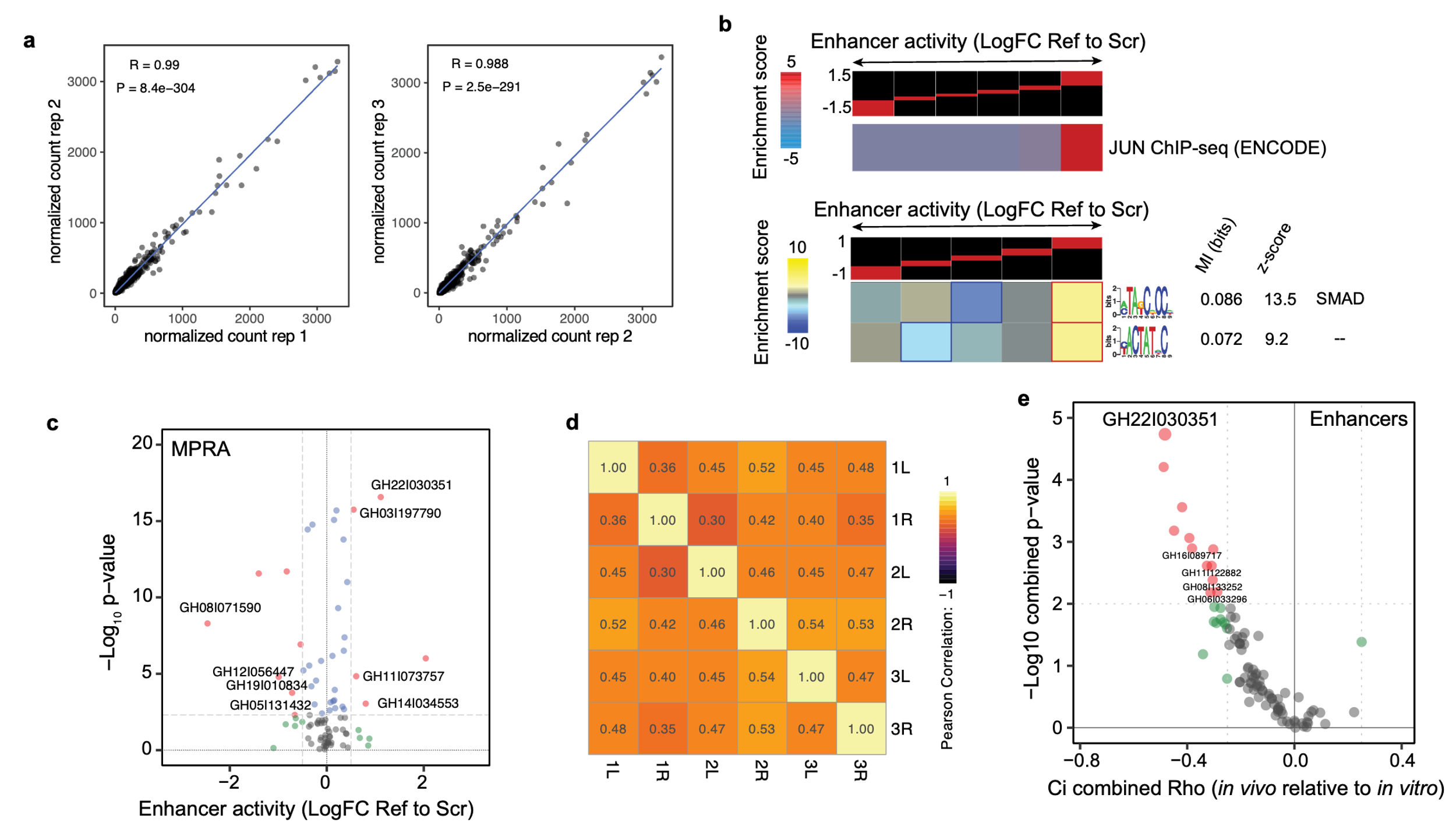

##### Figure S2. Functional characterization of hyper-mutated regulatory regions in prostate cancer.

**(a)** Our MPRA assay was done in biological triplicate. Here, we show the pair-wise comparison of normalized counts between replicates. The counts are normalized by library size factor. **(b)** Gene-set enrichment analysis of enhancer activity as measured by our MRPA measurements. For this analysis, the ratio of reference allele to scrambled control was used to sort regulatory regions from repressive (left) to activating (right). The values were then grouped into equally populated bins. In case of annotated ENCODE binding sites, iPAGE was used to identify the trans factor whose binding sites are enriched at the two ends of this spectrum. As shown here, JUND binding was significantly associated with increased enhancer activity. The bottom heatmap shows a similar heatmap for the discovered motifs. For each motif, FIRE reports the mutual information value (MI) and the associated z-score. The motifs were compared against the database of known motifs using Tomtom (MEME suite). **(c)** Enhancer activity of hypermutated regulatory regions as an aggregate of their assayed segments in our MPRA measurements. See methods of the details of this aggregation. **(d)** We performed CRISPRi *in vivo* screens in three mice (1-3), two flanks (L and R) each. Here, the pairwise correlation coefficients of sgDNA counts in each tumor are shown. The counts were summed across these tumors and compared to an *in-vitro*-grown library to calculate a fitness score associated with each regulatory region. **(e)** Aggregate phenotypic scores for each hypermutated regulatory region assayed for *in vivo* tumor growth in the C4-2B background. For this analysis, results from individual sgRNA activities targeting the same regulatory regions were combined into a singular measure.

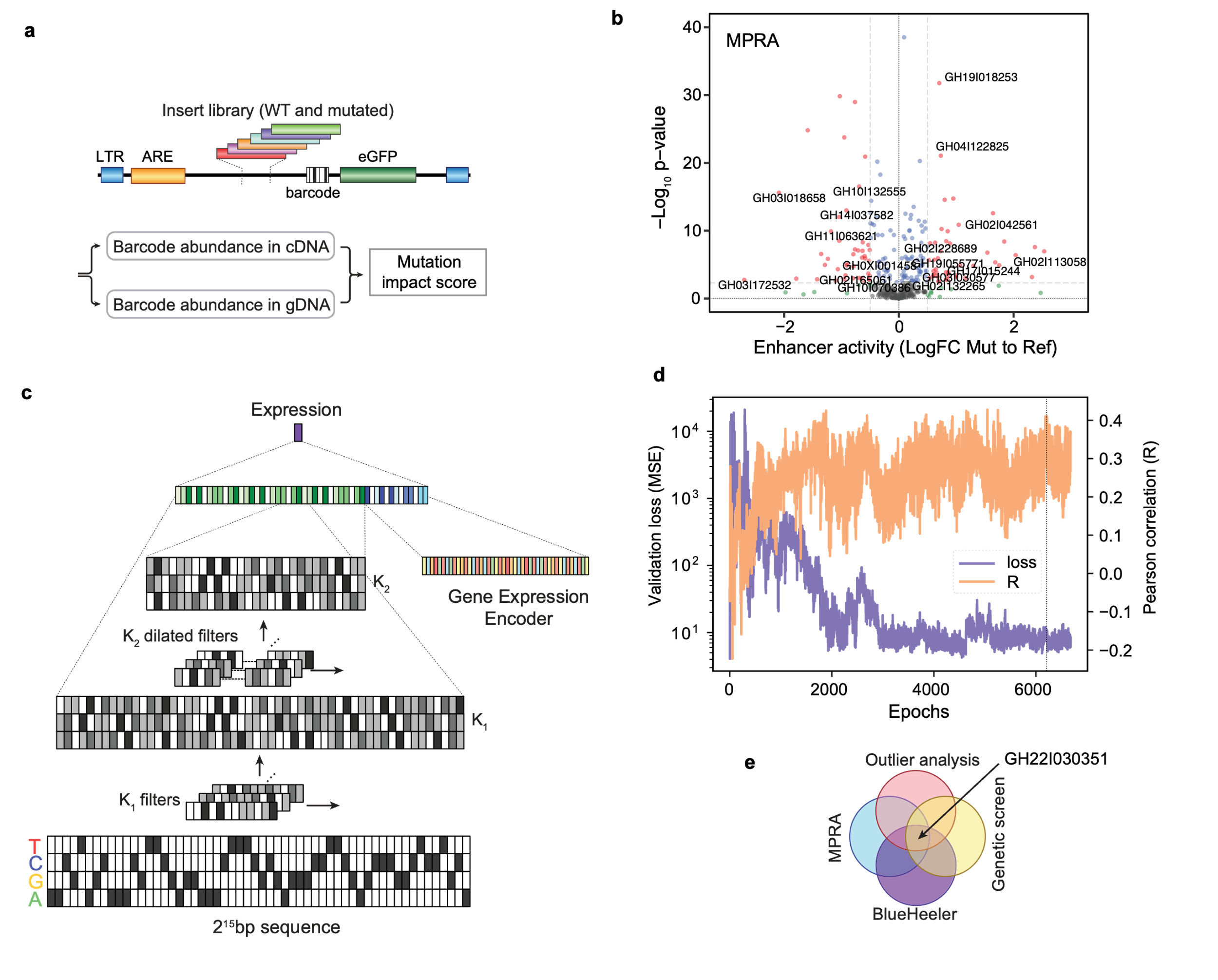

##### Figure S3. Measuring functional consequences of mutations in regulatory regions.

**(a)** Schematic of our MPRA setup for measuring the functional impact of mutations. **(b)** The combined effect of mutations in each regulatory region measured using MPRAs. **(c)** The general schematic of our Blue Heeler (BH) model, which combines sequence and cell-state embeddings to predict expression of a given gene. **(d)** The training of the BH model is over ~7,000 training batches. Shown here are the loss and Pearson correlation for the validation set. The final chosen model is marked by a dotted line. **(e)** Through a combined analysis of in culture, *in vivo*, and *in silico* observations, we nominated GH22I030351 as the strongest candidate non-coding driver in mCRPC.

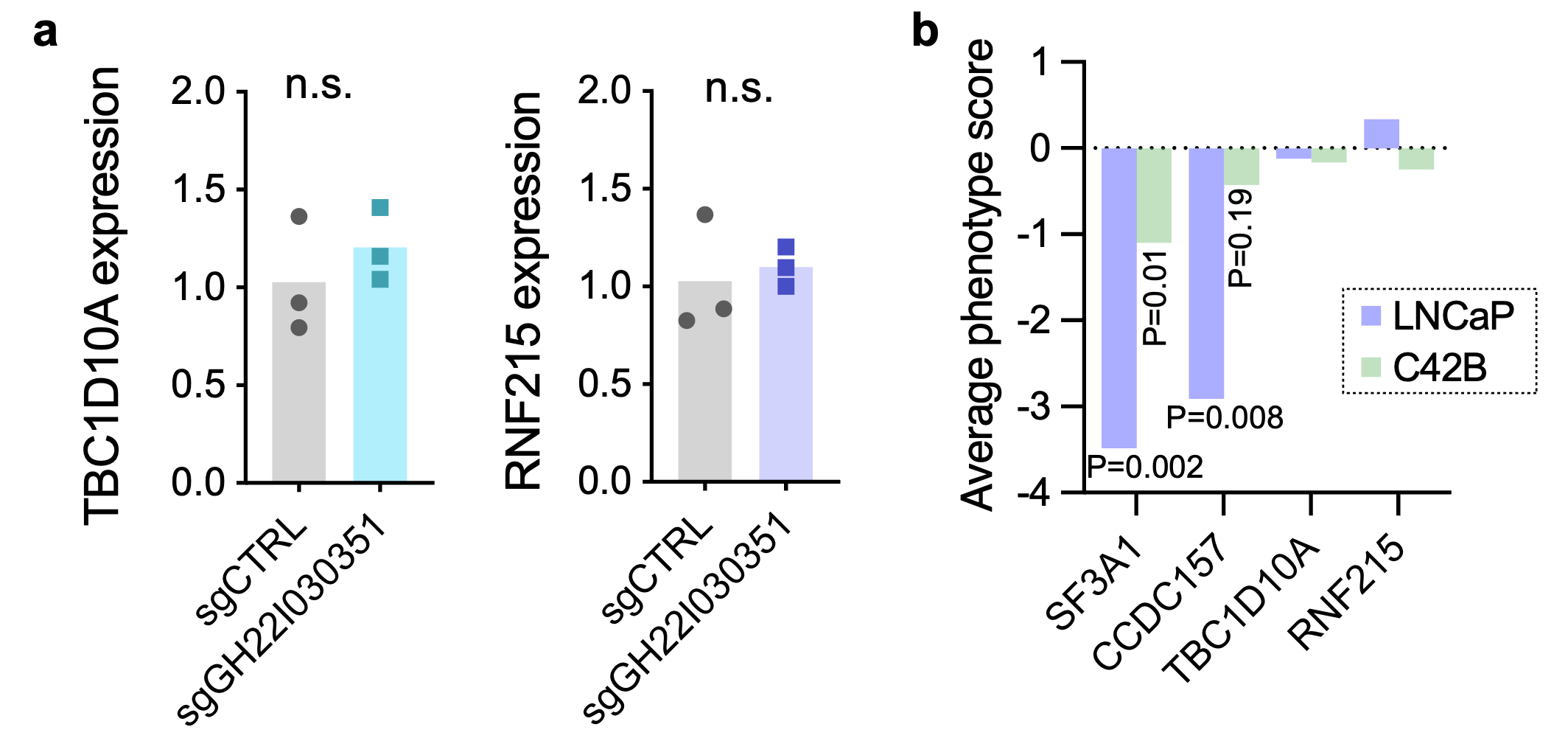

##### Figure S4. Functional targets of GH22I030351.

**(a)** Unlike SF3A1 and CCDC157, CRISPRi-mediated inhibition of GH22I030351 in C4-2B prostate cancer cells did not have an impact on the expression of TBC1D10A and RNF215. Therefore, the functional consequences of GH22I030351 silencing on prostate tumor growth in xenografts is unlikely to be through the function of these annotated target genes. **(b)** The phenotypic score associated with CRISPRi-mediated silencing of GH22I030351 downstream targets in two isogenic prostate cell lines, namely LNCaP and C4-2B, in a published large scale pooled *in vitro* growth screen (45). Negative scores imply reduced representation in the population upon knock-down.

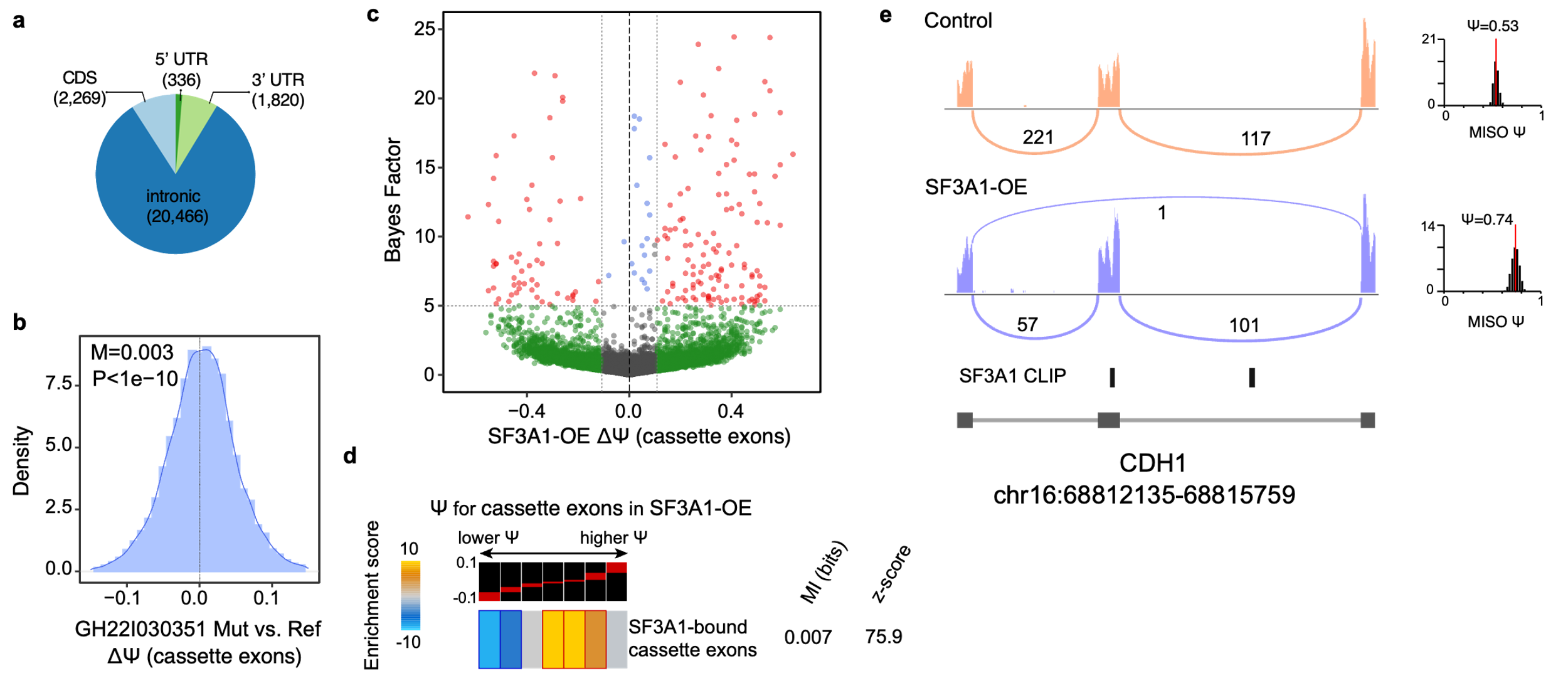

##### Figure S5. Splicing reprogramming through SF3A1 up-regulation.

**(a)** Annotation of SF3A1 binding sites, determined using CLIP-seq in the C4-2B prostate cancer cell line. As expected, the absolute majority of binding sites were in intronic regions. **(b)** Cassette exons that are bound by SF3A1 show a significant increase in their Ψ in GH22I030351-mutated samples in our mCRPC cohort. Reported are the median and Wilcoxon signed rank test. **(c)** Volcano plot of changes in alternative splicing patterns in cells over-expressing SF3A1 (C4-2B background). The analysis was performed using MISO and exons with ΔΨ >10% and Bayes factor >5 are marked as significant. **(d)** Enrichment of SF3A1-bound exons among those up-regulated upon SF3A1 over-expression, and their depletion among those with lower Ψ. Reported are the mutual information and the associated *z*-score. The ΔΨ bins with statistically significant enrichment or depletion (based on hyper-geometric *P*) are marked with a solid border. **(e)** A sashimi plot for exon 9 of CDH1 as an exemplary target of SF3A1. Shown here is the cassette exon, the identified SF3A1 binding sites, and the Ψ estimates for control and SF3A1 over-expression samples.

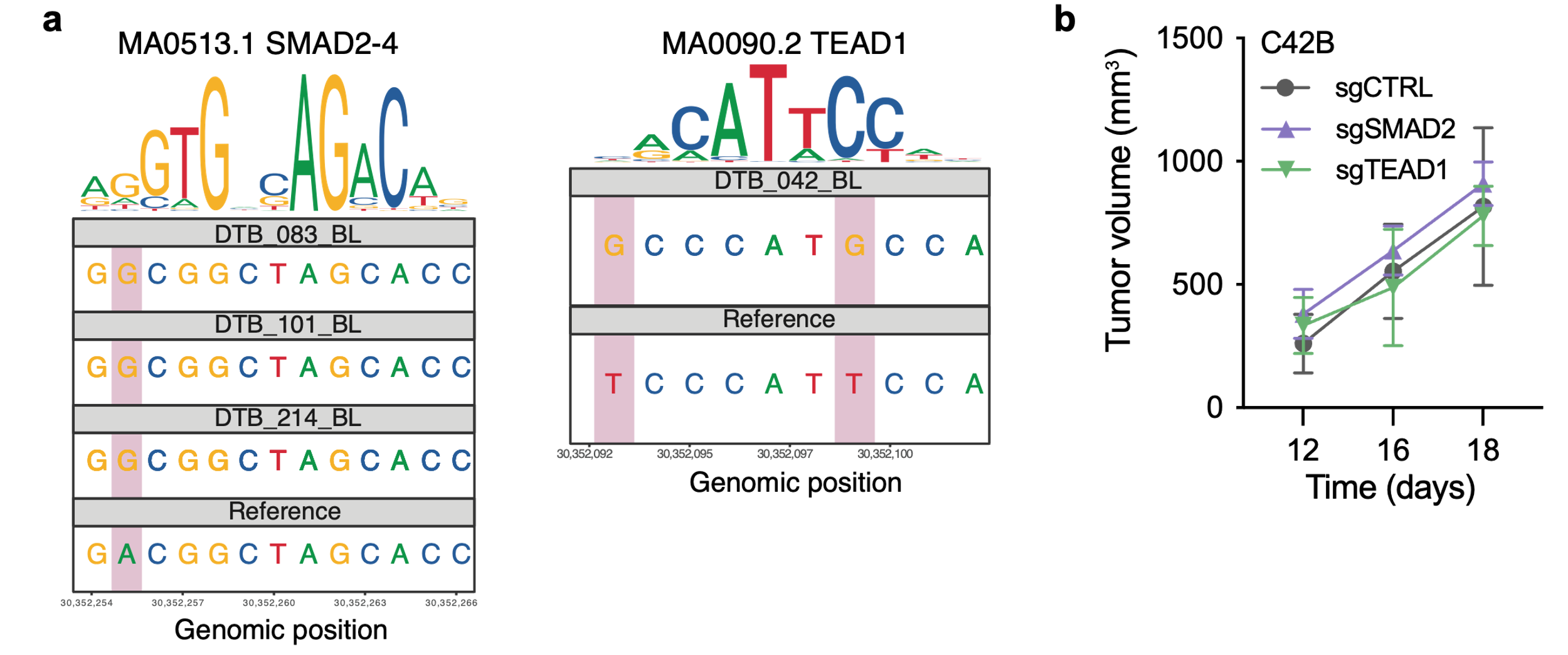

##### Figure S6. SOX6, SMAD2, and TEAD1 identified as putative transcription factors impacted by mutations in GH22I030351.

**(a)** Sequence motif with mutations observed among patients for SMAD2-4 (top) and TEAD 1 (bottom). The top part shows the sequence logo and the bottom panels correspond to a patient’s sequence or the reference GRCh38/hg38 genome sequence. **(b)** Unlike SOX6, silencing SMAD2 or TEAD1 did not significantly impact subcutaneous tumor growth in xenografted mice.
